# Microtubule Assembly and Pole Coalescence: Early Steps in *C. elegans* Oocyte Meiosis I Spindle Assembly

**DOI:** 10.1101/2020.02.03.933051

**Authors:** Chien-Hui Chuang, Aleesa J. Schlientz, Jie Yang, Bruce Bowerman

## Abstract

How oocytes assemble bipolar meiotic spindles in the absence of centrosomes as microtubule organizing centers remains poorly understood. We have used live cell imaging in *C. elegans* to investigate requirements for the nuclear lamina and for conserved regulators of microtubule dynamics during oocyte meiosis I spindle assembly, assessing these requirements with respect to recently identified spindle assembly steps. We show that the nuclear lamina is required for microtubule bundles to form a cage-like structure that appears shortly after oocyte nuclear envelope breakdown and surrounds the oocyte chromosomes, although bipolar spindles still assembled in its absence. Although two conserved regulators of microtubule nucleation, RAN-1 and γ-tubulin, are not required for bipolar spindle assembly, both contribute to normal levels of spindle-associated microtubules and spindle assembly dynamics. Finally, the XMAP215 ortholog ZYG-9 and the nearly identical minus-end directed kinesins KLP-15/16 are required for proper assembly of the early cage-like structure of microtubule bundles, and for early spindle pole foci to coalesce into a bipolar structure. Our results provide a framework for assigning molecular mechanisms to recently described steps in *C. elegans* oocyte meiosis I spindle assembly.

## INTRODUCTION

*C. elegans* oocyte meiotic spindles form in the absence of centrosomes and reduce the duplicated diploid genome to a haploid content through two rounds of cell division called meiosis I and II (Dumont and Desai, 2012; Mullen et al., 2019; Ohkura, 2015; Severson et al., 2016). While acentrosomal microtubule nucleation pathways have been shown to mediate meiotic and mitotic spindle assembly in some settings, how *C. elegans* oocytes nucleate microtubules and assemble bipolar spindles remains poorly understood. Conserved mechanisms for microtubule nucleation, including the small GTPase Ran pathway and γ-tubulin ring complexes, are not known to be required for oocyte meiotic cell division in *C. elegans* (Askjaer et al., 2002; McNally et al., 2006). Moreover, the augmin complex, which mediates microtubule branching in other animals, is not conserved in *C. elegans* (Edzuka et al., 2014). While microtubule severing by the conserved AAA+ ATPase complex called katanin promotes microtubule density in *C. elegans* oocytes, presumably by generating microtubule fragments that can further elongate (Srayko et al., 2006), mechanism(s) that nucleate microtubule substrates for katanin severing remain unknown.

Recent studies have defined a sequence of four steps that assemble bipolar spindles in *C. elegans* oocytes (Gigant et al., 2017; Mullen and Wignall, 2017) (Fig. 1A). First, oocyte nuclear envelope breakdown (NEBD) leads to a diffuse cloud of microtubule signal entering the nucleus. Second, microtubule bundles appear peripherally, underneath the dis-assembling nuclear lamina, to form a cage-like structure that surrounds the oocyte chromosomes. Third, the microtubule cage become organized such that multiple small foci of microtubules ends form in association with the pole-marker ASPM-1. Finally, these small pole foci coalesce to form a bipolar spindle as chromosomes congress to a metaphase plate (Connolly et al., 2015).

**Figure 1.**
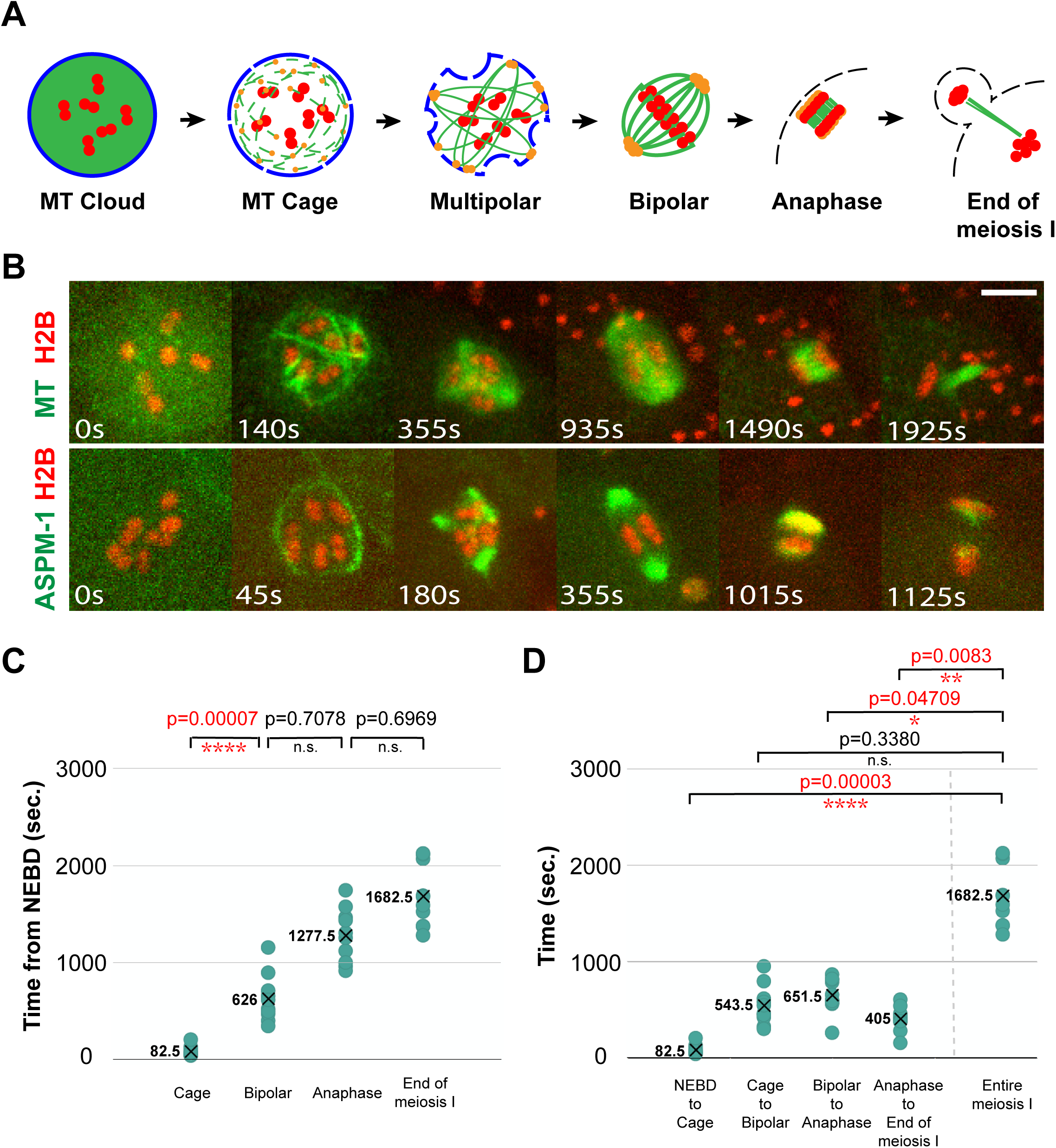
Wild-type oocyte meiosis I spindle assembly. (A) Schematic of *C. elegans* oocyte meiosis I spindle assembly. Nuclear lamina (blue), microtubules (MT, green), chromosomes (red), spindle poles (orange), and oocyte plasma membrane (black dashed line) are shown. (B) Time-lapse images (see Materials and Methods) of live control oocytes expressing either GFP::TBB-2 and mCherry::H2B to mark microtubules and chromosomes (upper row), or GFP::ASPM-1 and mCherry::H2B to mark spindle poles and chromosomes (lower row). Scale bars in all figures represent 5 μm. (C) Scatter plot showing the time from nuclear envelope breakdown (NEBD) to different stages of meiosis I in control oocytes expressing GFP::TBB-2 and mCherry::H2B. For all figures, the X and the adjacent number indicate the average value for each dataset, and variance was compared using the F-test to calculate p-values. (D) Scatter plot of the times from one stage to the next, and the entire time of meiosis I.

Defining these meiotic spindle assembly steps provides a foundation for identifying molecular mechanisms that participate in this reductive cell division. While the early microtubule cage assembles in close proximity to the nuclear lamina, how these microtubules are nucleated and whether the nuclear lamina is required for their assembly are not known. RNAi-mediated knockdown of two nearly identical minus-end directed *C. elegans* kinesin-14 family members, KLP-15 and −16, destabilizes the cage-like structure (Mullen and Wignall, 2017), consistent with the demonstrated ability of kinesin-14 family members to bundle parallel microtubules (Fink et al., 2009). The small GTPase Ran has been implicated in chromosome-mediated nucleation of microtubule assembly during meiosis and mitosis in some organisms (Clarke and Zhang, 2008), but not in *C. elegans* (Askjaer et al., 2002). RNAi knockdown of the *C. elegans* γ-tubulin TBG-1 does not prevent oocyte meiotic cell division, but TBG-1 is diffusely associated with oocyte meiotic spindles and its knockdown exacerbates the microtubule loss caused by reduced katanin function (McNally et al., 2006). Finally, the multiple TOG domain protein and XMAP215 family member in *C. elegans*, called ZYG-9, promotes astral microtubule stability during early embryonic mitosis and is known to be required for oocyte meiotic spindle assembly (Bellanger et al., 2007; Bellanger and Gonczy, 2003; Yang et al., 2003). XMAP215 orthologs act as microtubule polymerases in cooperation with γ-tubulin ring complexes (Gunzelmann et al., 2018; Thawani et al., 2018), and have roles in promoting both microtubule stability and instability (Kosco et al., 2001; Shirasu-Hiza et al., 2003), but the role of ZYG-9 in *C. elegans* oocytes remains poorly understood.

To improve our understanding of *C. elegans* oocyte meiosis I spindle assembly, we have used RNAi and mutations to reduce the function of several proteins implicated in this process, and live cell imaging with fluorescent protein fusions to assess their requirements. Here we report our analysis of requirements for the single nuclear lamina protein in *C. elegans* LMN-1 (Liu et al., 2000), the small GTPase RAN-1, the γ-tubulin TBG-1, the kinesin-14 family members KLP-15/16, and the XMAP215 ortholog ZYG-9. Our results indicate that the nuclear lamina is required for assembly of the cage-like array of KLP-15/16-stabilized microtubule bundles observed early in spindle formation, although bipolar spindles still assembled after LMN-1 knockdown. Furthermore, while not by themselves required for oocyte meiotic cell division, reduction of either RAN-1 or TBG-1 decreased spindle microtubule levels and altered spindle assembly dynamics.

Finally, we show that KLP-15/16 and ZYG-9 make distinct contributions both to assembly of the early microtubule cage structure and to the coalescence of pole foci to form a bipolar spindle.

## RESULTS

### Wild-type oocyte meiosis I spindle assembly

To observe early steps in oocyte meiosis I spindle assembly, we imaged control oocytes *in utero* using spinning disk confocal microscopy and simultaneous two-color live imaging (see Materials and Methods), first with a transgenic strain expressing within the germline a GFP fusion to β-tubulin (GFP::TBB-2) to mark microtubules and an mCherry fusion to a histone (mCherry::H2B) to mark chromosomes (Figure 1B and Supplemental Figure 1). To assess progression through meiosis I, we designated the time at which the value for histone intensity in the nucleoplasm became equal to the value for the cytoplasm, which marks the initiation of nuclear envelope breakdown (NEBD), as t = zero and collected z-stacks encompassing the oocyte volume every 5 seconds. In all 10 control oocytes, we observed the previously described steps in spindle assembly: the rapid appearance of GFP::TBB-2 signal around chromosomes; the assembly of peripheral microtubule bundles to form a cage-like structure surrounding the chromosomes; the appearance of a multipolar spindle during prometaphase; and the coalescence of multiple small pole foci to form a bipolar spindle by metaphase. Subsequently, during anaphase, most pole microtubules disappeared and central spindle microtubules assembled as the segregating chromosomes moved apart. Finally, we designated the time point with the lowest level of GFP::TBB-2 signal in the spindle area, prior to meiosis II, as the end of meiosis I.

To further characterize spindle assembly, we imaged live control oocytes using a GFP fusion to the pole-marker ASPM-1 (GFP::ASPM-1) and the mCherry::H2B fusion (Figure 1B, Supplemental Figure 2). As described previously (Gigant et al., 2017; Mullen and Wignall, 2017), GFP::ASPM-1 initially localized along microtubules during cage assembly but then became restricted to multiple small pole foci that coalesced into two poles by metaphase. These GFP::ASPM-1-marked poles subsequently broadened and faded during anaphase.

While all control oocytes in the GFP::TBB-2 background underwent a similar progression in spindle morphology, the time required to complete meiosis I varied from a minimum of 1280 seconds to a maximum of 2125 seconds, a 66% increase in duration. To determine when this variability arises, we assessed the variance in timing during progression through the different stages of assembly. The most significant increase in variance occurred during the transition from the appearance of the microtubule cage structure to the establishment of spindle bipolarity (Figures 1C and 1D). The subsequent transitions, from the establishment of bipolarity to anaphase onset, and from anaphase onset to the completion of meiosis I, did not show significant further increases. Having collected data sets that established a baseline for wild-type assembly dynamics, we next used RNAi to assess genetic requirements for the early spindle assembly steps and for timely progression through oocyte meiosis I.

### The nuclear lamina is required for assembly of the peripheral microtubule cage structure

Because the microtubule cage that assembles after the initiation of NEBD is adjacent to the nuclear lamina (Mullen and Wignall, 2017), we first asked if the nuclear lamina is required for cage assembly. After using RNAi in the GFP::TBB-2, mCherry::H2B background to knock down the single *C. elegans* lamin LMN-1, we did not detect any peripheral microtubule bundles in 8 of 10 oocytes, and in 2 oocytes we detected only a few relatively short bundles (Figure 2A, Supplemental Figure 3). Although the cage was missing or greatly reduced after LMN-1 knockdown, the overall level of microtubule signal was similar to that observed in control oocytes (Figure 2C), and the subsequent steps in spindle assembly appeared relatively normal, with transient multi-polar structures that coalesced to form bipolar spindles of normal pole-to-pole length (Figure 2B and 2D). Moreover, we still observed the disappearance of most microtubules from the poles as central spindle microtubules appeared during anaphase between the segregating chromosome masses, but the extent of chromosome segregation was reduced compared to control oocytes, and in 5 of 20 oocytes, imaged using either GFP::TBB-2 or GFP::ASPM-1, chromosome segregation completely failed (Figure 2E).

**Figure 2.**
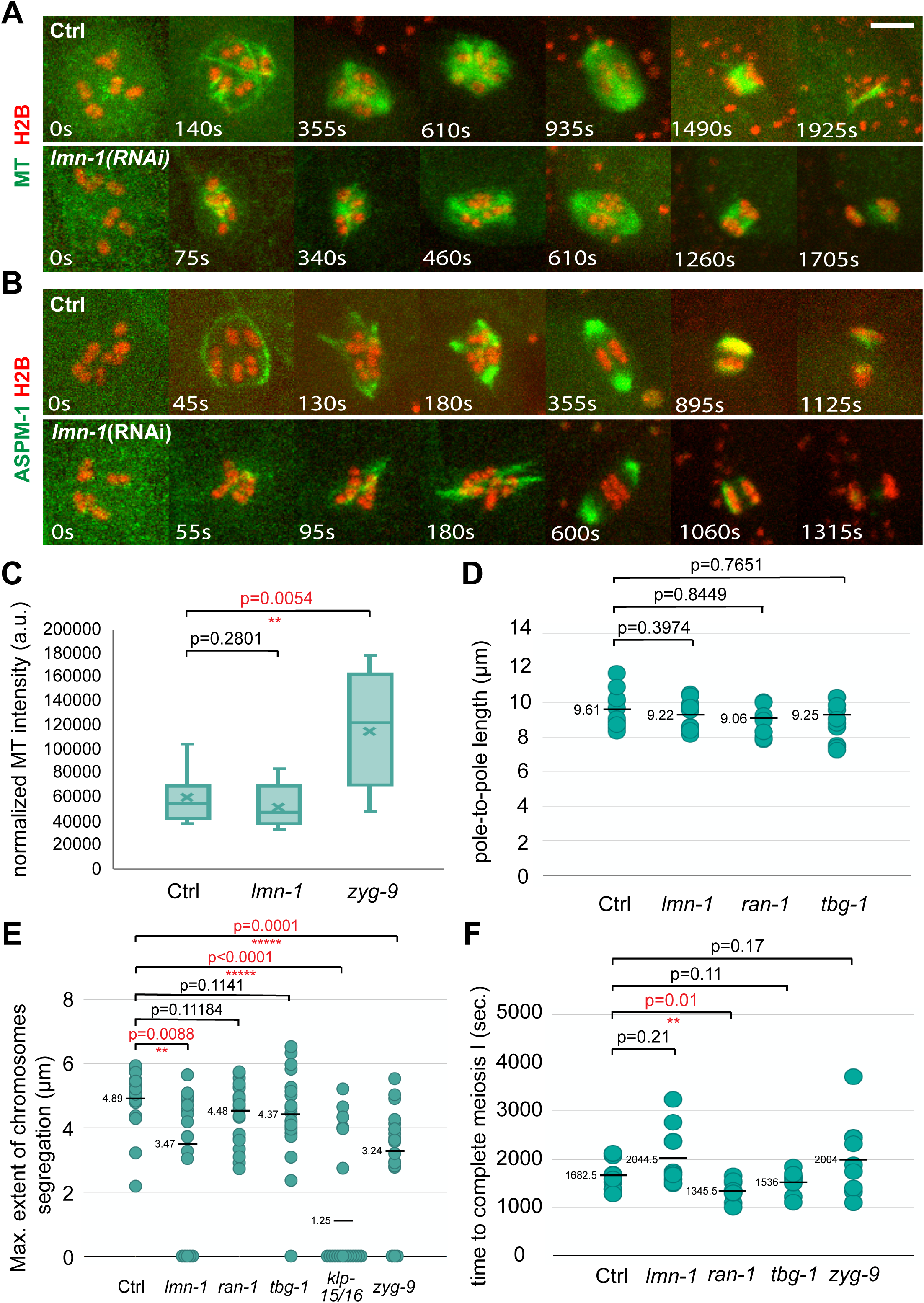
The nuclear lamina is required for the early cage-like microtubule structure during oocyte meiosis I. (A-B) Time-lapse images during meiosis I of live control and *lmn-1(RNAi)* oocytes expressing either GFP::TBB-2 and mCherry::H2B (A), or GFP::ASPM-1 and mCherry::H2B (B). (C) Normalized microtubule pixel intensity in arbitrary units. The boxplot displays the datasets and the median (line) and mean (x) values for the 3 continuous time points with the highest pixel intensities for control and mutant oocytes. (D) Pole-to-pole length measured at metaphase I in control and mutant oocytes expressing GFP::ASPM-1 and mCherry::H2B. For all figures, distributions of scatter plot values were compared using the Mann-Whitney U test to calculate p-values. (E) Scatter plot showing maximum extent of chromosome segregation at the end of meiosis I for control and mutant oocytes expressing either GFP::TBB-2 and mCherry::H2B or GFP::ASPM-1 and mCherry::H2B. Numbers of oocytes that failed to segregate chromosomes at the end of meiosis I (i.e. maximum extent of chromosome segregation value = 0) were: 5 for *lmn-1(RNAi)*, 1for *tbg-1(RNAi)*, 12 for *klp-15/16(RNAi)* and 3 for *zyg-9(RNAi)* oocytes. (F) Scatter plot showing the time required to progress through meiosis I in control and mutant oocytes expressing GFP::TBB-2 and mCherry::H2B.

Using the pole marker GFP::ASPM-1 (Figure 2B, Supplemental Figure 4), cage-like microtubule bundles were again greatly reduced or absent, and as in control oocytes, GFP::ASPM-1 was detected along microtubules as multipolar spindles assembled. However, the GFP signal persisted along microtubules for a longer period of time before concentrating at pole foci as spindle bipolarity was established. Finally, despite these altered spindle dynamics, the time required to progress through meiosis I, although more variable, was not on average significantly different compared to control oocytes (Figure 2F). We conclude that the nuclear lamina is required for assembly of the microtubule cage that surrounds chromosomes early in meiosis I. While the cage structure is not required for assembly of a bipolar spindle, the dynamics of spindle assembly were altered and chromosome segregation was in some cases entirely absent after LMN-1 knockdown.

### RAN-1 promotes microtubule assembly and normal oocyte nuclear size

We next asked if known regulators of microtubule nucleation might also be required for assembly of the microtubule bundles that form the early cage structure. Previous studies of the *C. elegans* small GTPase RAN-1 and the γ-tubulin family member TBG-1 have not found them to be required for oocyte meiotic cell division (see Introduction). However, the early cage structure is not essential for bipolar spindle assembly after LMN-1 knockdown, and whether cage assembly occurs in the absence of RAN-1 or TBG-1, and the impact of these regulators on spindle assembly dynamics, have not been addressed.

We used RNAi to knock down RAN-1 in strains expressing GFP::TBB-2 and mCherry::H2B (Figure 3A and Supplemental Figure 6), GFP::ASPM-1 and mCherry::H2B (Figure 3B and Supplemental Figure 7), and GFP::LMN-1 and mCherry::TBB-2 (Figures 3C). We observed a roughly normal sequence of spindle morphology transitions, with a microtubule and ASPM-1 cage forming soon after NEBD, followed by the appearance of multiple small pole foci that coalesced to form a bipolar spindle of normal length that segregated chromosomes to a normal extent (Figures 2D, 2E, 3A and 3B, Supplemental Figures 6 and 7). However, compared to control oocytes, the quantities of microtubules detected throughout meiosis I were reduced (Figure 3D), and the microtubule cage was smaller in diameter (Figure 3A and 3B, Supplemental Figures 6, 7 and 8A). Furthermore, the time required to complete meiosis I, based on GFP::TBB-2 imaging (Figure 2F), was reduced from an average of 1682.5 seconds in control oocytes to an average of 1345.5 seconds after RAN-1 knockdown (p = 0.01). We further assessed progression through meiosis 1 based on chromosome dynamics and observed a significant decrease during the time from NEBD to metaphase (Figure 3E). We also examined spindle pole dynamics with GFP::ASPM-1 and again found that the decrease in time occurred during the establishment of spindle bipolarity, between NEBD and metaphase (Supplemental Figure 5A). To summarize, RAN-1 is at least partially required for spindle microtubule assembly during meiosis I but may not be required for a functional bipolar spindle to form. Surprisingly, reducing RAN-1 function leads to a more rapid progression through meiosis I that may result from a reduction in the time required for early pole foci to coalesce and form a bipolar spindle.

**Figure 3.**
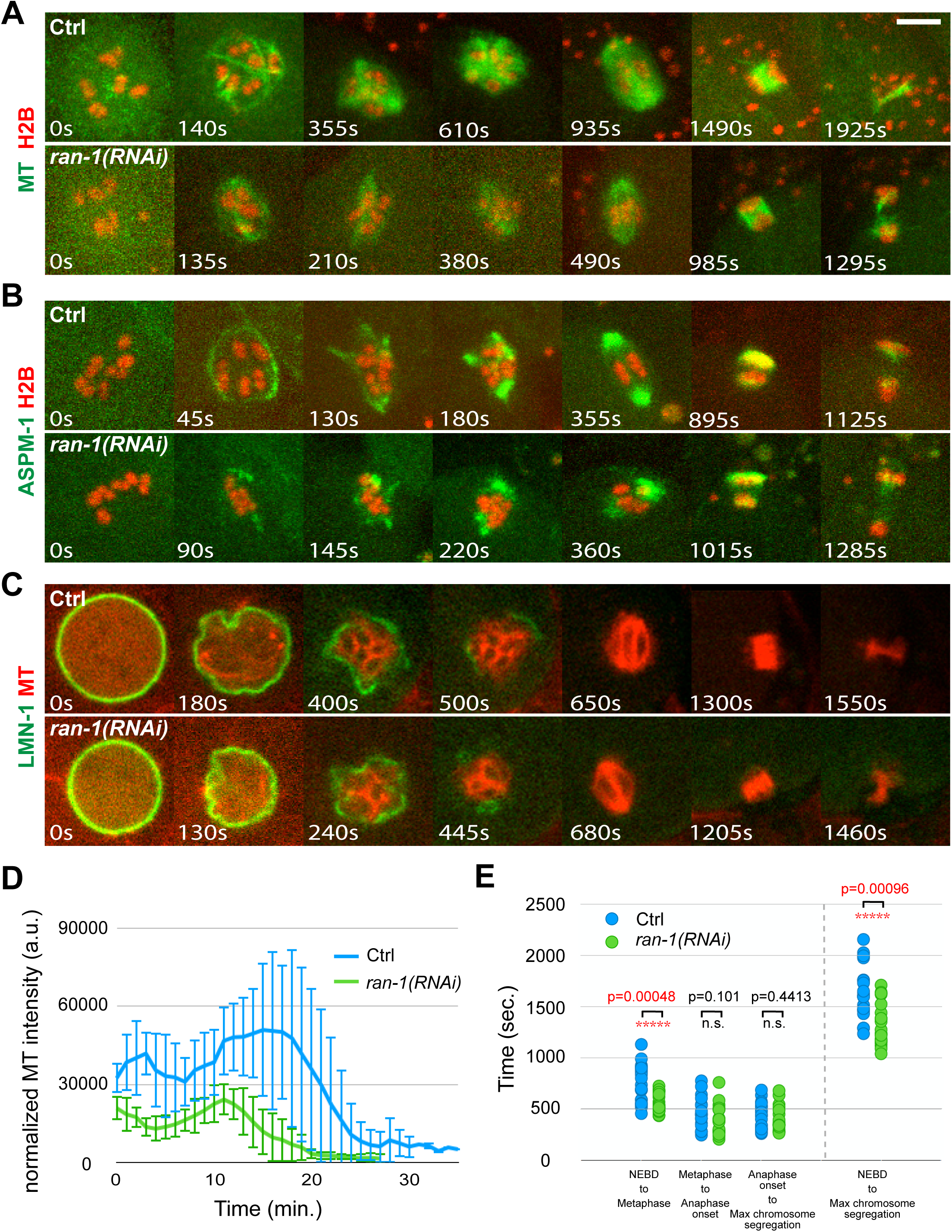
RAN knockdown reduces oocyte nuclear size and microtubule levels during oocyte meiosis I. (A-C) Time-lapse images during meiosis I for live control and *ran-1(RNAi)* oocytes expressing GFP::TBB-2 and mCherry::H2B (A), GFP::ASPM-1 and mCherry::H2B (B), or GFP::LMN-1 and mCherry::TBB-2 to mark the nuclear lamina and microtubules (C). Maximum intensity z-projections (A-B), or the middle plane of the nucleus (C) are shown. (D) Normalized microtubule pixel intensity in arbitrary units measured over time with 1-minute time intervals. Time 0 = NEBD. For all figures, error bars depict one standard deviation at each time point. (E) Comparison of the length of time from one stage to the next between the control and *ran-1(RNAi)* oocytes from strains expressing either GFP::TBB-2 and mCherry::H2B or GFP::ASPM-1 and mCherry::H2B. NEBD: the time at which the value for histone intensity in the nucleoplasm became equal to the value for the cytoplasm; Metaphase: time at which chromosomes aligned on the metaphase plate; Anaphase onset: the time at which chromosomes started to separate; Maximum chromosome segregation: the time at which chromosomes separated to the maximum distance.

**Figure 6.**
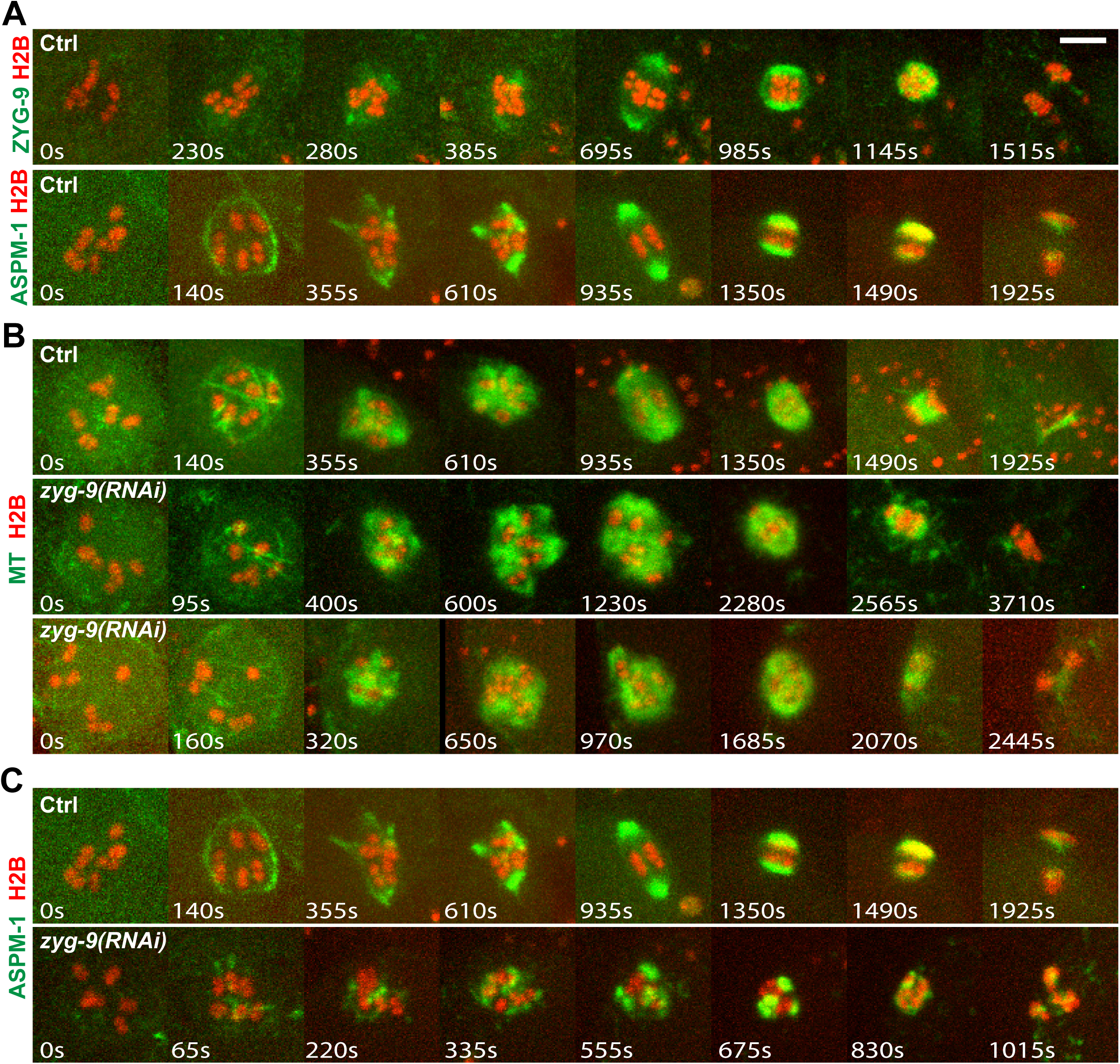
ZYG-9 knockdown causes spindle assembly defects throughout oocyte meiosis I. (A) Time-lapse images during meiosis I of live control oocytes expressing either GFP::ZYG-9 and mCherry::H2B (upper row), or GFP::ASPM-1 and mCherry::H2B (lower row). (B-C) Time-lapse images during meiosis I of live control and *zyg-9(RNAi)* oocytes expressing either GFP::TBB-2 and mCherry::H2B (B), or GFP::ASPM-1 and mCherry::H2B (C).

**Figure 7.**
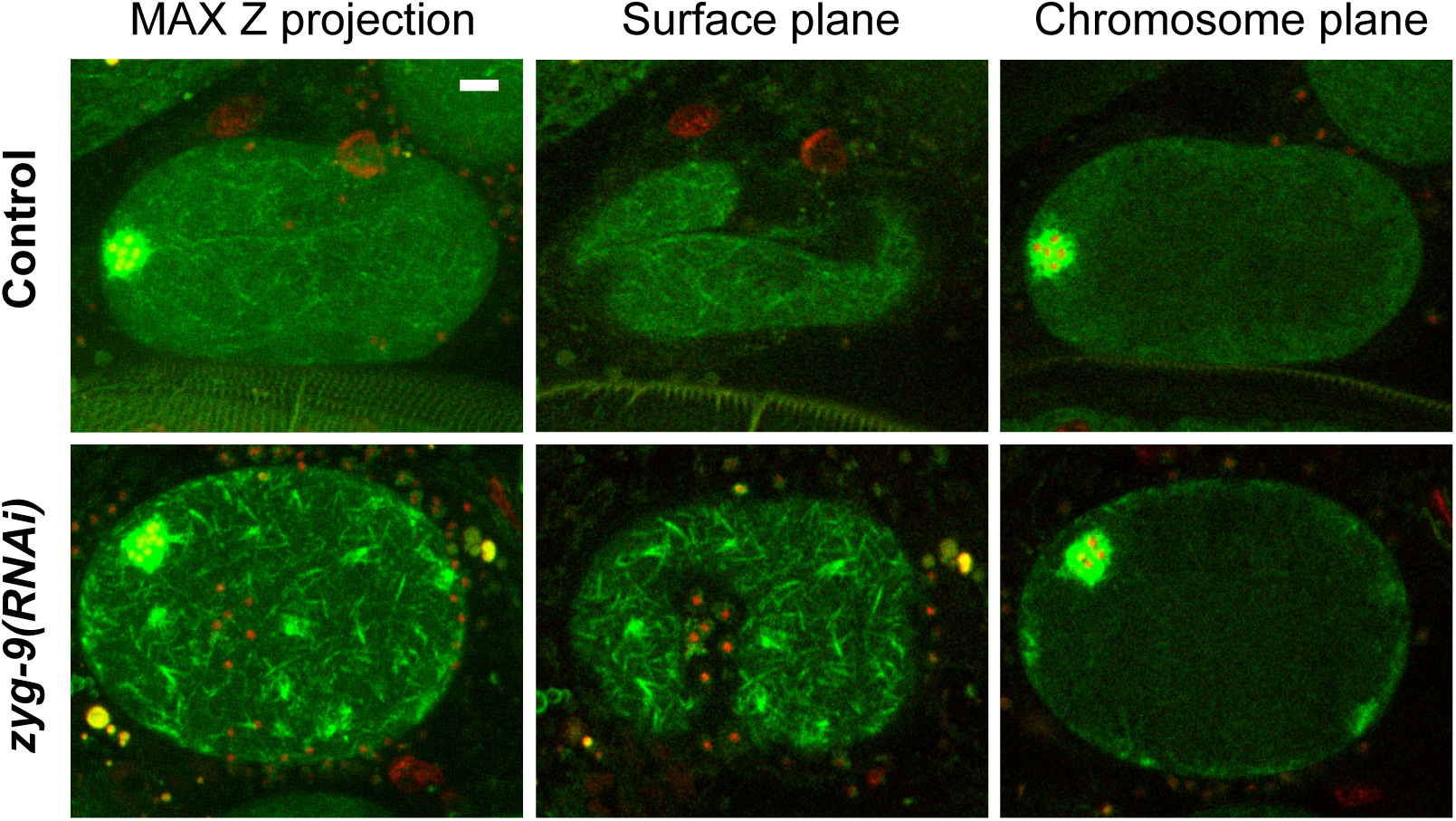
Microtubule levels are increased throughout the oocyte cortex during meiosis I after ZYG-9 knockdown. *Ex utero* spinning disk confocal images for live control (upper row) and *zyg-9(RNAi)* (lower row) oocytes expressing GFP::TBB-2 and mCherry::H2B. Pixel intensity was enhanced to in the green channel to highlight cortical microtubules. Left panels: maximum intensity z-projection image of 15 planes with 1 μm z-spacing. Middle panels: surface plane. Right panels: chromosome plane.

Finally, we asked if the smaller microtubule cage structure observed after RAN-1 knockdown is associated with a smaller oocyte nucleus, or if the microtubule cage forms more internally relative to the nuclear lamina in normally sized nuclei. To address this issue, we knocked down RAN-1 in a transgenic strain expressing GFP::LMN-1 and an mCherry fusion to TBB-2 (Figure 3C). While the timing of NEBD, based on the fragmentation and decrease of the GFP::LMN-1 signal over time appeared normal, the diameters of the oocyte nuclei were reduced to roughly the same extent as the microtubule cage structure (Figure 3A-3C and Supplemental Figures 8A and 8B). Similarly, the diameters of oocyte nuclei measured using Nomarski optics were reduced relative to control oocytes (Supplemental Figure 8C and 8D). Consistent with these findings, RAN-1 knockdown has previously been reported to result in the production of fertilized embryos with abnormally small oocyte pronuclei (Askjaer et al., 2002).

### TBG-1 is required for proper oocyte nuclear positioning and promotes both microtubule levels and normal spindle assembly dynamics

We next examined spindle assembly after knocking down TBG-1 in transgenic strains expressing GFP::TBB-2 and mCherry::H2B (Figure 4A and Supplemental Figure 9), or GFP::ASPM-1 and mCherry::H2B (Figure 4B and Supplemental Figure 10). Prior to NEBD and spindle assembly, we observed a displacement of oocyte nuclei from the cortex (Fig. 4C), followed by a reduction in microtubule levels throughout meiosis I (Fig. 4D). The diameter of the microtubule cage structure, though more variable, was on average not significantly different from its diameter in control oocytes (Supplemental Figures 8A), but the subsequent dynamics of spindle assembly were distinct from those observed in both control and RAN-1 knockdown oocytes. Shortly after cage assembly, the microtubules collapsed into a small cluster around the oocyte chromosomes, rather than forming the peripheral multipolar structure observed in control oocytes. Similarly, GFP::ASPM-1 foci and the oocyte chromosomes became grouped together in a tight cluster. Subsequently, more extended microtubule structures appeared and multiple GFP::ASPM-1 foci emerged peripherally to the oocyte chromosomes and coalesced to form a bipolar spindle of normal length that segregated chromosomes to a normal though more variable extent (Figures 2D and 2E). In one case, when imaging GFP::ASPM-1, we observed a complete failure in chromosome segregation (Figure 2E). Finally, the time to complete meiosis I and the variance in the time required to achieve spindle bipolarity were similar to control oocytes (Figures 1C, 2F and 4E). In summary, as reported previously (McNally et al., 2006), TBG-1 was not required for bipolar spindle assembly. However, microtubule levels were reduced and the dynamics of spindle assembly were altered and distinct from those observed after RAN-1 knockdown.

**Figure 4.**
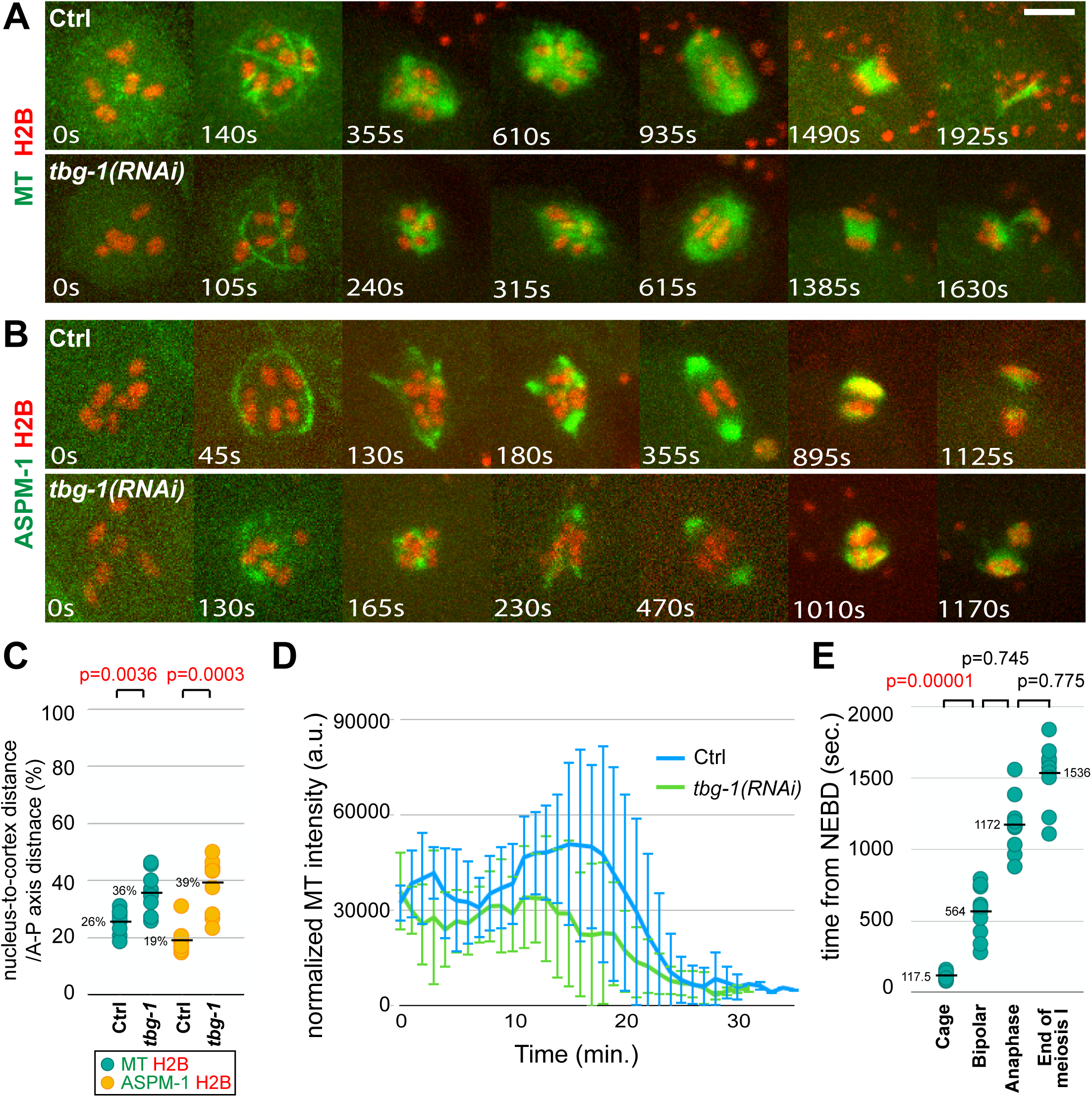
TBG-1 knockdown alters spindle dynamics after the microtubule cage-like structure appears and reduces microtubule levels during oocyte meiosis I. (A-B) Time-lapse images during meiosis I of live control and *tbg-1(RNAi)* oocytes expressing either GFP::TBB-2 and mCherry::H2B (A), or GFP::ASPM-1 and mCherry::H2B (B). (C) Scatter plot showing nuclear position for control and *tbg-1(RNAi)* oocytes, measured at NEBD as the distance from the center of the nucleus to the closest edge of the oocyte on the same focal plane/the length of oocyte anterior-posterior axis. (D) Normalized microtubule pixel intensity in arbitrary units measured over time with 1-minute time intervals. Time 0 = NEBD. (E) Scatter plot showing the time from NEBD to different stages of meiosis I in *tbg-1(RNAi)* oocytes expressing GFP::TBB-2 and mCherry::H2B.

### KLP-15 and −16 are required for cage stability and promote pole coalescence

The nearly identical *C. elegans* minus-end directed kinesin-14 family members KLP-15 and −16 have been shown previously to have a role in stabilizing the microtubule bundles that form the early cage structure (Mullen and Wignall, 2017). After RNAi knockdown of KLP-15/16, the bundles were less prominent compared to control oocytes and a more diffuse spherical distribution of microtubules then surrounded the chromosomes, which underwent segregation in only about half of the depleted oocytes. As these results were obtained using RNAi, and spindle structures were assessed using immunofluorescence in fixed oocytes, we have analyzed KLP-15/16 requirements using live cell imaging with putative null alleles.

We first made a strain carrying likely null alleles for both *klp-15* and *-16*, which are on the same chromosome separated by about 3.5 map units (Figure 5A). Using CRISPR/Cas9, we introduced a small deletion near the 5’ end of the first coding exon of *klp-16* in a strain homozygous for a previously isolated *klp-15* deletion allele, *klp-15(ok1958)*. The CRISPR-generated *klp-16* deletion resulted in a frameshift followed by multiple stop codons and likely eliminates all gene function. The resulting double mutant chromosome, *klp-15(ok1958) klp-16(or1952)*, was then balanced with a marked inversion chromosome, *tmc18* (Dejima et al., 2018). When homozygous, these two mutations resulted in the development of fertile adults with reduced brood sizes and highly penetrant and recessive embryonic lethality (Figure 5D).

**Figure 5.**
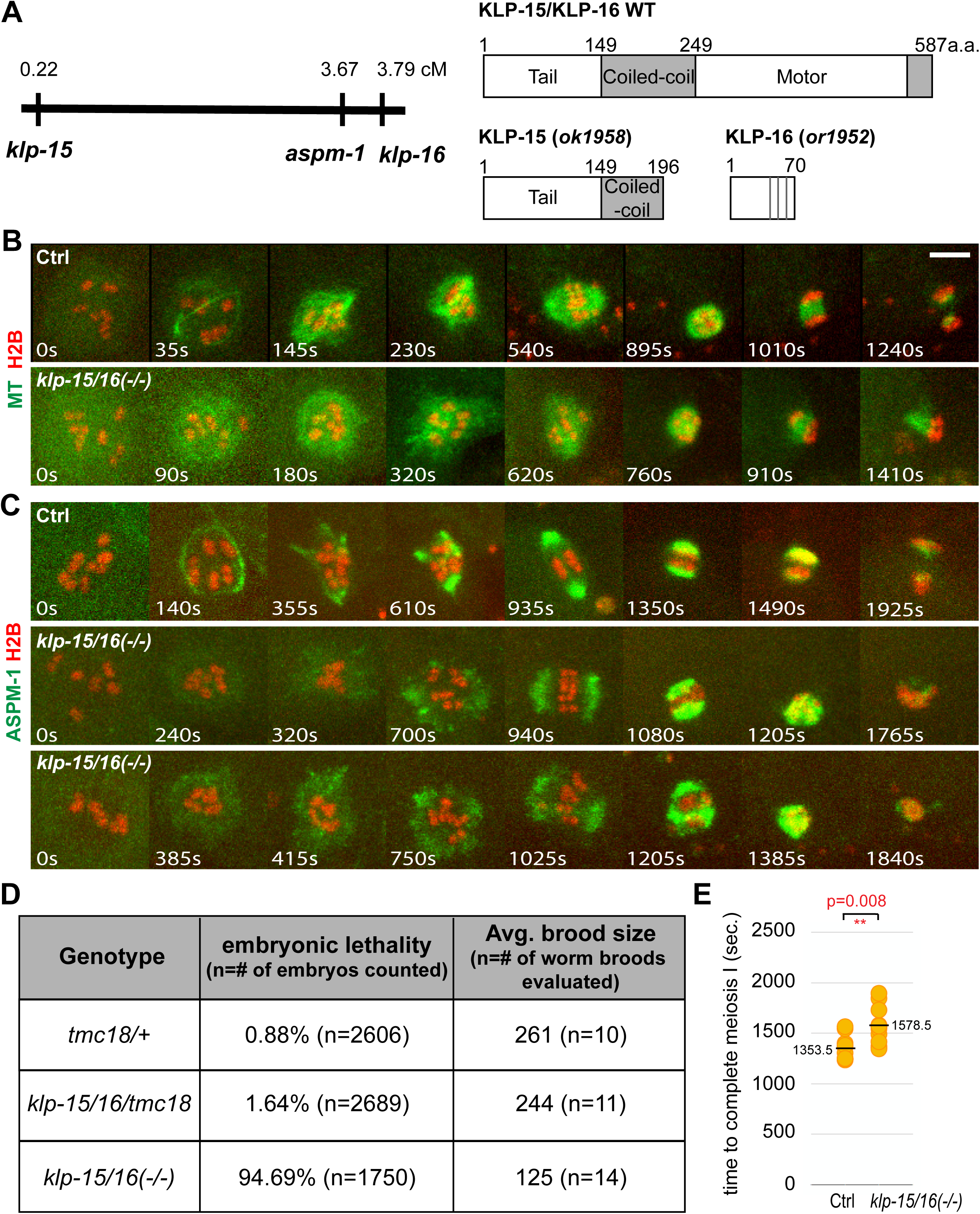
KLP-15/16 knockdown destabilizes cage-like microtubule bundles and reduces pole coalescence. (A) Left panel: locations of *klp-15*, *aspm-1* and *klp-16* on chromosome I. Right panel: domain architectures of wild-type KLP-15/16 and the KLP-15/16 double mutant used in this study. Vertical grey lines indicate stop codons that follow the frameshift caused by the *or1952* deletion. (B) Time-lapse images during meiosis I of control and *klp-15/16(-/-)* oocytes expressing GFP::TBB-2 and mCherry::H2B. (C) Time-lapse images during meiosis 1 of control or *klp-15/16(-/-)* double mutant oocytes expressing GFP::ASPM-1and mCherry::H2B. (D) Embryonic lethality and average brood sizes for *klp-15/16(-/-)* double mutant and control strains. (E) Scatter plot showing the time required to progress through meiosis I for control and *klp-15/16(-/-)* double mutant oocytes.

We next used live imaging to examine oocyte meiosis I spindle assembly in the *klp-15/16* null mutant background. We used genetic crosses to generate a *tmc18* balanced *klp-15/16* double mutant strain that expresses GFP::TBB-2 and mCherry::H2B (Figure 5B and Supplemental Figure 11). Because the *aspm-1* gene resides on the same chromosome between *klp-15* and *-16* (Figure 5A), we obtained a CRISPR/Cas9-generated *in situ* GFP fusion to the endogenous *aspm-1* locus on the *klp-15(or1958) klp-16(or1952)* chromosome (see Materials and Methods), and used genetic crosses to introduce the mCherry::H2B fusion (Figure 5C and Supplemental Figure 12). We observed a decreased prominence of the microtubule bundles that form the cage, as reported previously (Mullen and Wignall, 2017), and a subsequent failure to form a bipolar spindle or segregate chromosomes in over half of the mutant oocytes (Figures 2E and 5C, Supplemental Figures 11 and 12). Live imaging of the *klp-15/16* double mutant with GFP::ASPM-1 also suggested that the microtubule cage was reduced in prominence. Subsequently, GFP::ASPM-1 foci were detected both peripheral to and amongst the chromosomes (Figure 5C, Supplemental Figure 12 and Supplemental Movies 1-3). These GFP::ASPM-1 foci became more prominent over time, often forming broad poles that failed to coalesce into the more compact structures observed in control oocytes (Figure 1B and Supplemental Figure 2). In some cases, broad arrays of GFP::ASPM-1 foci nearly encircled the chromosomes, and the average time to complete meiosis I was increased (Figure 5E). We conclude that in addition to being important for stable assembly of the microtubule cage and chromosome segregation, KLP-15/16 also are important for the coalescence of early pole foci into properly organized spindle poles.

### ZYG-9 restricts microtubule bundles to the periphery during cage assembly and promotes pole coalescence and stability

The XMAP215 family member ZYG-9 is known to be required for oocyte meiotic spindle assembly (Yang et al., 2003), but the nature of this requirement remains poorly understood. To examine its role, we first assessed the dynamics of ZYG-9 localization in comparison to microtubules and ASPM-1. We used CRISPR/Cas9 to generate an *in situ* fusion of GFP to the endogenous *zyg-9* locus and genetic crosses to introduce the mCherry::H2B fusion (Figure 6A and Supplemental Figure 15A). In contrast to GFP::ASPM-1, we did not detect GFP::ZYG-9 in association with the microtubule cage; rather it was initially more diffusely present near chromosomes and subsequently became enriched at multiple pole foci and also was present more diffusely throughout the spindle during pole coalescence. Upon the establishment of spindle bipolarity, GFP::ZYG-9 was enriched at the poles but in contrast to GFP::ASPM-1, GFP::ZYG-9 also was detected between the poles. ZYG-9 binds the coiled-coil TACC ortholog TAC-1, and both promote microtubule stability during early embryonic mitosis in *C. elegans* (Bellanger et al., 2007; Bellanger and Gonczy, 2003). We therefore used CRISPR/Cas9 to generate an *in situ* fusion of GFP to the endogenous *tac-1* locus and observed localization dynamics similar to GFP::ZYG-9 (Supplemental Figure 15B). To summarize, ZYG-9 and its partner TAC-1 were both present throughout spindle assembly and both appeared more restricted in distribution than microtubules but, as assembly progressed, more broadly distributed than ASPM-1.

We next examined ZYG-9 requirements after RNAi knockdown in transgenic strains expressing GFP or mCherry fusions to TBB-2, ASPM-1 and H2B and observed multiple defects during meiosis I spindle assembly. First, when imaging spindles marked with GFP::TBB-2 or GFP::ASPM-1, microtubule bundles assembled to form a peripheral cage shortly after NEBD, but some of the microtubule bundles were not restricted to the periphery and instead passed through the interior of the chromosome occupied space (Figure 6B and 6C, Supplemental Figures 13 and 14, and Supplemental Movies 4-9). Subsequently, foci of GFP::TBB-2 and GFP::ASPM-1 were observed not only at the periphery surrounding oocyte chromosomes, but also between some of the bivalent chromosomes. Moreover, small spindle-like structures often appeared to form around individual or small groups of bivalents, in contrast to the peripheral spindle foci that coalesced to form a bipolar spindle in control oocytes.

The abnormal dynamics of spindle assembly observed after ZYG-9 knockdown were accompanied by a significant increase in the level of oocyte spindle microtubules (Figures 2C and 6B, Supplemental Figure 13; Supplemental Movies 10 and 11), and a variable but significant increase in the diameter of the microtubule cage structure (Supplemental Figure 8A). We also observed increased microtubule levels throughout the oocyte cortex during meiosis I (Figure 7). These increases were surprising because ZYG-9 is required for the stability of astral microtubules during early embryonic mitosis (Bellanger et al., 2007; Bellanger and Gonczy, 2003), and ZYG-9 orthologs in other species promote microtubule assembly (Akhmanova and Steinmetz, 2015), although in some contexts they also promote microtubule instability (Shirasu-Hiza et al., 2003).

We also observed a striking lack of pole stability and extensive chromosome segregation defects after ZYG-9 knockdown. In control oocytes, early small pole foci stably associated with each other over time (Figure 1B, Supplemental Figures 2 and 16, Supplemental Movies 1 and 12). In contrast, after ZYG-9 knockdown, pole foci marked by GFP::ASPM-1 fused and then often broke apart as meiosis I progressed (Figure 6C, Supplemental Figure 16 and Supplemental Movies 13-15). Consistent with a role in pole stability, chromosomes sometimes segregated into three masses during anaphase (10 of 20 oocytes), although in other cases no segregation (3 of 20 oocytes) or segregation into two masses (7 of 20 oocytes) were observed (Supplemental Figures 5C, 13 and 14). Finally, we observed similar defects throughout oocyte meiosis I after knocking down the TAC-1 binding partner for ZYG-9 (Supplemental Movies 16 and 17), indicating that ZYG-9 and TAC-1 have similar if not identical requirements. In summary, ZYG-9 and TAC-1 restrict cage microtubule bundles to the periphery and promote pole coalescence and stability. Notably, they also appear to limit both spindle and cortical microtubule levels during oocyte meiosis I.

## DISCUSSION

We have examined the requirements for factors involved in *C. elegans* oocyte meiotic spindle assembly, with the goal of assessing their roles during a sequence of four recently described assembly steps that generate these acentrosomal and yet bipolar spindles during meiosis I. Our results show that the nuclear lamina, comprised of the single *C. elegans* nuclear lamin LMN-1, is required for assembly of the underlying cage-like microtubule structure, although this structure is not required for bipolar spindle assembly. While knockdown of either of two conserved regulators of microtubule nucleation, the small GTPase RAN-1 and TBG-1/γ-tubulin, did not prevent bipolar spindle assembly or chromosome segregation, microtubule levels after both knockdowns were reduced and spindle assembly dynamics were altered. We also identified two additional contributions to assembly of the early cage-like network of microtubule bundles. After knockdown of the XMAP215 ortholog ZYG-9, or its binding partner TAC-1, the microtubule bundles were no longer restricted to the periphery but in some cases passed through the space occupied by oocyte chromosomes. As reported previously, these microtubule bundles were reduced in prominence in mutants lacking the nearly identical minus-end directed kinesins KLP-15 and −16. Finally, both ZYG-9 and KLP-15/16 were required for early spindle pole foci to coalesce into a bipolar structure, but in distinct ways. ZYG-9 knockdown resulted in a lack of pole stability during coalescence, while pole foci failed to coalesce in *klp-15/16* mutant oocytes. Our results, considered in more detail below, document requirements for microtubule nucleation, cage assembly and pole coalescence, key steps in the assembly of acentrosomal oocyte meiosis I spindles in *C. elegans*.

### Microtubule nucleation and C. elegans oocyte meiotic spindle assembly

Although several genes are known to be required for proper organization of oocyte meiotic spindles in *C. elegans*, how microtubules are nucleated during this process has remained poorly understood (Severson et al., 2016). Two widely conserved regulators that contribute to microtubule nucleation in other contexts, γ-tubulin and the small GTPase Ran, have not been found to be required for oocyte meiotic cell division in *C. elegans* (see Introduction). However, our results indicate that both contribute to producing normal microtubule levels and spindle assembly dynamics.

RAN-1 knockdown did not appear to substantially affect any of the steps in oocyte spindle assembly: the early microtubule cage formed, small pole foci then appeared and coalesced into a bipolar spindle of normal length that segregated chromosomes to the same extent as in control oocytes. However, RAN-1 knockdown did result in the production of oocyte nuclei that were smaller in diameter, and the microtubule cage also was reduced in diameter. Moreover, microtubule levels declined to a minimum earlier than was observed in control oocytes, and the time required to progress through meiosis I was reduced.

TBG-1 knockdown also did not prevent assembly of the microtubule cage, and its diameter was similar to those in control oocytes. However, rather than proceeding to form a network of peripherally located small spindle pole foci, the cage collapsed into a condensed ball of microtubule signal surrounding the chromosomes. Subsequently, microtubules emerged from the collapsed structure, and the pole marker ASPM-1 appeared in multiple foci that coalesced to form a bipolar spindle of normal length that segregated chromosomes to the same extent as in control oocytes. In spite of these changes in spindle assembly dynamics, we did not detect any defects in chromosome segregation by the end of meiosis I after knockdown of either RAN-1 or TBG-1, with one exception after TBG-1 knockdown, consistent with previous reports indicating the lack of an essential requirement for either of these regulators during oocyte meiosis.

While the more normal sequence of assembly events after RAN-1 knockdown and the more substantially altered assembly dynamics after TBG-1 knockdown suggest that these two regulators play distinct roles, in neither case have we been able to determine the consequences of fully eliminating gene function. For both RAN-1 and TBG-1, we analyzed oocyte meiotic defects at a time prior to which the RNAi treatments resulted in adult sterility (Materials and Methods), indicating that their activities were not entirely absent. More complete elimination of their functions could result in more severe and perhaps more similar defects. Indeed, defining gene requirements for oocyte meiotic spindle assembly is challenging because essential genes involved in this process are often required for fertile adults to develop. Due to earlier requirements, null alleles often result in zygotic embryonic or larval lethality, or adult sterility, precluding their use for more definitively analyzing requirements during oocyte meiosis. A recently developed alternative approach to reducing gene function at different times in development is to use CRISPR/Cas9 to tag endogenous loci with a degron motif that induces degradation of the tagged protein upon treatment with the plant hormone auxin (Zhang et al., 2015). While this approach does not allow one to conclusively determine null phenotypes, it can be especially useful for simultaneously reducing the functions of multiple degron-tagged proteins, as RNAi often requires different time courses for depleting different loci and is less reliable as more genes are simultaneously targeted. Degron tagging therefore may facilitate better assessment of the potentially distinct roles of RAN-1 and TBG-1.

### The nuclear lamina as a platform for assembling the cage-like network of microtubule bundles

While we did not detect a requirement for either RAN-1 or TBG-1 in assembly of the early microtubule cage, we did find that the nuclear lamina, which directly overlies this structure, is required for its assembly. RNAi knockdown of the only *C. elegans* lamin LMN-1 nearly eliminated the peripheral microtubule bundles. Restricting the assembly of early microtubule bundles to the periphery to form a cage-like network might promote pole coalescence by having it occur only along the inner surface of the nuclear lamina, rather than throughout the volume occupied by oocyte chromosomes. Consistent with such a role, the time required to complete meiosis I in control oocytes differs due to variability in the time required for pole coalescence, and the reduced time required to complete meiosis after RAN-1 knockdown occurs during pole coalescence and correlates with a reduction in the diameter and surface area of the cage-like network.

We also detected later abnormalities as spindle assembly progressed after LMN-1 knockdown. The pole marker ASPM-1 appeared to persist along the length of microtubules for a longer period of time compared to control oocytes, before becoming enriched at the two spindle poles. Bipolar spindles of normal length ultimately assembled and the time required to complete meiosis I was more variable but not significantly increased compared to control oocytes. While alternative bipolar spindle assembly mechanisms appear to compensate for loss of the microtubule cage structure, chromosomes were segregated to a lesser extent, and in 5 of 20 cases chromosome segregation failed. Some or all of these defects could be indirectly due to disruptions in the nuclear import of factors required for a fully functional spindle to assemble, and we did detect variable and sometimes lower levels of some kinetochore proteins after LMN-1 knockdown (data not shown). Nevertheless, our results indicate that the nuclear lamina plays an important role in *C. elegans* oocyte meiotic spindle assembly, and roles for lamins in spindle assembly have also been reported in *Xenopus* extracts and during *Drosophila* male meiosis(Goodman et al., 2010; Hayashi et al., 2016; Tsai et al., 2006).

### ZYG-9/XMAP215 and the kinesin-14 family members KLP-15/16 contribute to proper assembly of the cage-like network of microtubule bundles

In addition to the nuclear lamina being required for assembly of the microtubule cage early in meiosis I spindle assembly, we also found requirements for ZYG-9 and the minus-end directed kinesins KLP-15/16 in the assembly of this structure. As reported previously based on RNAi knockdown (Mullen and Wignall, 2017), we found that the cage microtubule bundles that formed in oocytes from worms homozygous for likely null alleles of *klp-15* and *-16* were less prominent and rapidly became undetectable, with the microtubules instead forming a diffuse cloud encompassing the oocyte chromosomes, and chromosome segregation often completely failed.

We observed a very different defect in the microtubule cage structure after depletion of ZYG-9/XMAP215, or of its binding partner TAC-1. Stable and prominent microtubule bundles formed peripherally to the oocyte chromosomes early in spindle assembly, as in control oocytes, but some of the bundles passed through the interior of the chromosome occupied volume, rather than being restricted to the periphery. Subsequently, pole foci formed not only at the periphery but also amongst the chromosomes, and oocytes often failed to assemble bipolar spindles. In some oocytes, the spindles became tripolar and segregated chromosomes into three distinct masses, while in other cases a bipolar structure failed to emerge and chromosome segregation completely failed. We conclude that ZYG-9 and KLP-15/16 make distinct contributions to assembly of the cage-like network of microtubule bundles: KLP-15/16 are required for their stability, while ZYG-9 appears to restrict their assembly to the periphery.

In addition to the abnormal spatial organization of microtubule bundles and early spindle pole foci after ZYG-9 knockdown, we also observed increased levels both of spindle-associated microtubules and of microtubules throughout the oocyte cortex during meiosis I. This was surprising given that ZYG-9 orthologs have been reported to promote microtubule stability and act as microtubule polymerases (Akhmanova and Steinmetz, 2015). Indeed, astral microtubules during early embryonic mitosis in *C. elegans zyg-9* mutants are abnormally short (Bellanger et al., 2007; Bellanger and Gonczy, 2003). Nevertheless, other studies also have reported a role for XMAP215 orthologs in promoting instability (Brittle and Ohkura, 2005; Shirasu-Hiza et al., 2003), and our analysis of ZYG-9 provides further evidence that these TOG domain proteins can promote microtubule instability. Moreover, ZYG-9 can promote both microtubule stability and microtubule instability depending on the cellular context, even when the different activities are closely spaced in time, as also appears to be true in budding yeast (Kosco et al., 2001; Shirasu-Hiza et al., 2003). Finally, our results indicate that ZYG-9 acts in concert with its conserved binding partner, TAC-1, not only to promote microtubule stability during early embryonic mitosis, but also to promote microtubule instability during oocyte meiosis I.

A role for ZYG-9 in promoting microtubule instability may account for the spatial organization of the microtubule bundles that surround the oocyte chromosomes early in meiosis I spindle assembly. GFP::ZYG-9 and GFP::TAC-1 initially were distributed diffusely throughout the space occupied by chromosomes and were not detected in association with the peripheral microtubule bundles. Thus ZYG-9 and TAC-1 might prevent microtubule assembly in the volume occupied by chromosomes, restricting cage formation to the periphery.

### ZYG-9/XMAP215 and the kinesin-14 family members KLP-15/16 make distinct contributions to pole coalescence

In addition to being required for proper assembly of the microtubule cage, ZYG-9 and KLP-15/16 also appear to make distinct contributions to pole coalescence. In control oocytes, multiple small GFP::ASPM-1 pole foci moved toward each other and fused upon coming into contact, only rarely undergoing fission into distinct foci after merging. How these foci move toward each other remains unknown, but a similar process of coalescence has been observed in mouse oocytes (Schuh and Ellenberg, 2007). After ZYG-9 knockdown, we observed frequent examples of pole instability, in which pole foci would merge but then split apart. This process continued for an extended period of time with spindles frequently failing to become bipolar and often segregating chromosomes into three masses instead of two, or entirely failing to segregate chromosomes. By contrast, in *klp-15/16* mutant oocytes, pole foci became more prominent over time but were less dynamic. Although these foci often formed bipolar structures, the poles were much broader, and in some cases they nearly encircled the chromosomes without ever forming distinct poles. In addition, chromosome segregation often failed entirely. We conclude that while KLP-15/16 promote pole coalescence, ZYG-9 promotes pole stability, with both playing important roles in establishing a bipolar spindle.

Our results do not provide direct mechanistic insight into how either ZYG-9 or KLP-15/16 promote pole stability and coalescence, but the known functions of these conserved proteins may be relevant. Mutations in the *Drosophila* ortholog of KLP-15/16, Ncd, also result in disorganized oocyte meiotic spindles, with unfocused poles in some cases (Matthies et al., 1996; Skold et al., 2005), and Ncd also promotes pole assembly in acentrosomal *Drosophila* S2 cells (Goshima et al., 2005; Ito and Goshima, 2015), suggesting that these minus-end directed kinesins may have conserved roles in pole assembly. Like KLP-15/16, Ncd is localized throughout oocyte meiosis I spindles (Hatsumi and Endow, 1992; Mullen and Wignall, 2017), and the ability of kinesin-14 family members to cross-link parallel microtubules might contribute to pole coalescence (Fink et al., 2009).

With respect to the pole instability caused by loss of ZYG-9 or TAC-1, both also promote microtubule instability during oocyte meiosis I, raising the possibility that excessive microtubule growth might disrupt pole coalescence. In *Drosophila*, the ZYG-9 and TAC-1 orthologs Minispindles and D-TACC also are enriched at oocyte meiotic spindle poles, and loss of their function often results in tripolar spindles (Cullen and Ohkura, 2001), although the dynamics of pole stability have not been reported. Moreover, *Drosophila* Ncd is required for Minispindles to localize to oocyte meiotic spindle poles, and thus this minus-end directed kinesin-14 family member has been proposed to transport Minispindles to oocyte spindle poles and thereby promote pole assembly.

Finally, ZYG-9 can promote mitotic spindle assembly *in vitro* when incorporated into a centrosomal matrix that undergoes phase transitions (Woodruff et al., 2017), and the mouse XMAP215 and TACC orthologs chTOG and TACC3 have been reported to undergo phase transitions during mouse oocyte meiotic spindle assembly (So et al., 2019). Given the broad distribution of ZYG-9 and its binding partner TAC-1 during *C. elegans* oocyte meiotic spindle assembly, this protein complex may undergo phase transitions that influence microtubule and spindle pole stability. Future studies that assess in more detail the dynamics of ZYG-9/TAC-1 during oocyte meiosis I, and how ZYG-9/TAC-1 and KLP-15/16 interact, should further advance our understanding of how these regulators contribute to the assembly of bipolar but acentrosomal oocyte meiotic spindles.

## Materials and Methods

### C. elegans strains

*C. elegans* strains used in this study are listed in Supplemental table 1. All strains were maintained at 20°C on standard nematode growth medium plates seeded with *E. coli* strain OP50.

### RNAi

All RNAi experiments were carried out by feeding *E. coli* strain HT115(DE3) induced to express double-stranded RNA corresponding to each gene as previously described (Kamath et al., 2001; Timmons and Fire, 1998). The bacteria clones were picked from an RNAi library (Kamath et al., 2003). Synchronized larvae were washed with M9 three times and then plated on the induced plates and grown at 20°C until imaging. The feeding time varies from gene to gene in order to achieve maximum gene reduction without causing sterilization. For *tac-1*, worms were fed for 96 to 100 hours; for *tbg-1* and *zyg-9*, 48 to 52 hours; for *lmn-1*, 24 to 28 hours and for *ran-1*, 16 to 20 hours.

### CRISPR

#### Generation of *gfp::zyg-9* and *gfp::tac-1* transgenic strains

The appropriate sgRNA and PAM sites for *zyg-9* were selected by using the website http://crispr.mit.edu/. The repair oligo for *gfp::zyg-9* was obtained by asymmetric PCR using primers containing flanking bases at both the 5’ and 3’ ends of the PAM site to amplify the GFP-coding region from pCFJ150-GFP(dpiRNA)::CDK-1 (Zhang et al., 2018); Addgene plasmid#107938). The injection mixture of *gfp::zyg-9* repair oligo, co-CRISPR marker *dpy-10* repair oligo(Arribere et al., 2014, IDT), *dpy-10* crRNA(GCTACCATAGGCACCACGAG, IDT), trRNA (IDT) and Cas9-NLS nucleases (IDT) were injected into wild-type N2 young adults. The F1 progeny of the injected animals were selected for the roller phenotype and screened for GFP expression. The non-roller/dumpy F2 progeny of the F1 animals with correct GFP expression were identified and then further outcrossed with N2.

The transgenic strain of *gfp::tac-1* was made by the same approach as described above but with *gfp::tac-1* repair oligo and *tac-1* crRNA(CAACACAACCTTCACCAAAG, IDT).

#### Generation of double deletion strain of *klp-15* and *klp-16*

The *klp-15(ok1958) klp-16(or1952)* double mutant strain was generated by the same approach as described above, with a few modifications. The injection mix was injected into *klp-15(ok1958)* single mutants, with *klp-16* crRNA(TACTATCGGAGCACCGCCGA, IDT) and no repair template was provided. Injected hermaphrodites were kept at 15°C, and their broods were screened for *dpy-10* roller or dumpy co-conversion worms. Broods produced by hermaphrodites with the *dpy-10* co-conversion marker were screened for potential *klp-15/16* double mutant phenotypes (embryonic lethality), and lines identified as possibly carrying mutations to both *klp-15* and *klp-16* were balanced and Sanger sequenced after PCR amplification to identify the CRISPR/Cas9 induced mutation. The *in situ* fusion of GFP to the endogenous *aspm-1* locus in the *klp-15(ok1958) klp-16(or1952)* background was made by SunyBiotech using CRISPR/Cas-9 with the allele designation *GFP::aspm-1(syb1260)*.

### Image acquisition

*In utero* filming of oocytes was accomplished by mounting young adult worms with single row of embryos on a 5% agarose pad, with 1.5μl each of M9 buffer and 0.1μm polystyrene microspheres (Polysciences Inc.) on a microscope slide covered with a coverslip. *Ex utero* filming of oocytes for data in Supplemental Figure 16A and Supplemental Movies 10 and 11 was done by cutting open young adult worms in 3μl of egg buffer (118mM NaCl, 48mM KCl, 2mM CaCl_2_, 2mM MgCl_2_, and 25mM HEPES, PH 7.3) on a coverslip before mounting onto a 2% agarose pad on a microscope slide; all other data is from *in utero* imaging. Nomarski images were acquired on AxioSkop compound microscope (Zeiss) equipped with CCD camera using ImageJ software (National Institutes of Health). Fluorescence imaging was performed using a Leica DMi8 microscope outfitted with a spinning disk confocal unit – CSU-W1 (Yokogawa) with Borealis (Andor), dual iXon Ultra 897 (Andor) cameras, and a 100x HCX PL APO 1.4-0.70NA oil objective lens (Leica). Metamorph (Molecular Devices) imaging software was used for controlling image acquisition. For *in utero* movies of oocytes from GFP::LMN-1, mCherry::H2B and GFP::TAC-1, mCherry::H2B strains, the 488nm and 561nm channels were imaged simultaneously every 10 seconds with 1μm Z-spacing; every 5 seconds for oocytes from all other transgenic strains expressing fluorescent markers. 14 focal planes/z-stack were collected for all *klp-15/16* mutant oocytes and the control oocytes expressing GFP::TBB-2, mCherry::H2B in Figure 6; 21 focal planes/z-stack were collected for all other oocytes. Time lapse images in figures depict maximum projections for all fluorescent proteins except for GFP::LMN-1, which were single focal planes of maximum oocyte diameters. Time lapse supplemental movies show maximum projections for all z-stack focal planes unless otherwise indicated.

### Image processing and analysis

General imaging process—including merging red/green channels, cropping, stabilizing and z-projected images—was performed with ImageJ software (National Institutes of Health). Three-dimensional projection and rotation movies were made by using imaris software (Bitplane).

Normalized microtubule pixel intensity quantification was carried out by ImageJ software. The spindle area was determined by generating auto-threshold (set on Otsu) in a cropped area in the microtubule channel around the spindle in the projected z stack (Max intensity) treated with gaussian blur (sigma set at 2) at time points throughout meiosis I. The regions of interest (ROIs) were selected accordingly and saved in ROI manager. Measurements of microtubule intensity were taken by applying the saved ROIs to the projected z stack (sum slices) in the microtubule channel at the corresponding time points. Both the area of the ROIs and the mean gray value (MeanGV) were automatically calculated. In addition, an area excluding the spindle was selected and the MeanGV was calculated for the cytoplasm. Measurements of chromosome intensity were taken by selecting the oocyte chromosome area in the histone channel in the projected z stack (Max intensity) at the corresponding time points and the maximum gray value (MaxGV) were calculated. Additionally, an area excluding the oocyte and the sperm chromosomes was selected and the MaxGV was calculated for the cytoplasm. All of the measurements were placed into the following formula: (MeanGV (spindle)-MeanGV (cytoplasm))/(MaxGV (chromosomes)-MaxGV (cytoplasm)) × area(spindle) = normalized microtubule pixel intensity.

### Statistics

p-values comparing distributions for all scatter plots were calculated using the Mann-Whitney U test. p-values comparing variance for all scatter plots were calculated using the F-test.

## Acknowledgements

We thank The *Caenorhabditis elegans* Genetics Center (funded by the National Institutes of Health Office of Research Infrastructure Programs; P40 OK010440) for *C. elegans* strains, Chris Doe and Diana Libuda for sharing laboratory equipment, Adam Fries from the University of Oregon Imaging Core Facility for advice on microscope maintenance and use, and members of the Bowerman and Libuda laboratories for helpful discussions.

## Data Availability

All data sets used in the preparation of this manuscript are available upon request.

---

**Supplemental Figure 1.**
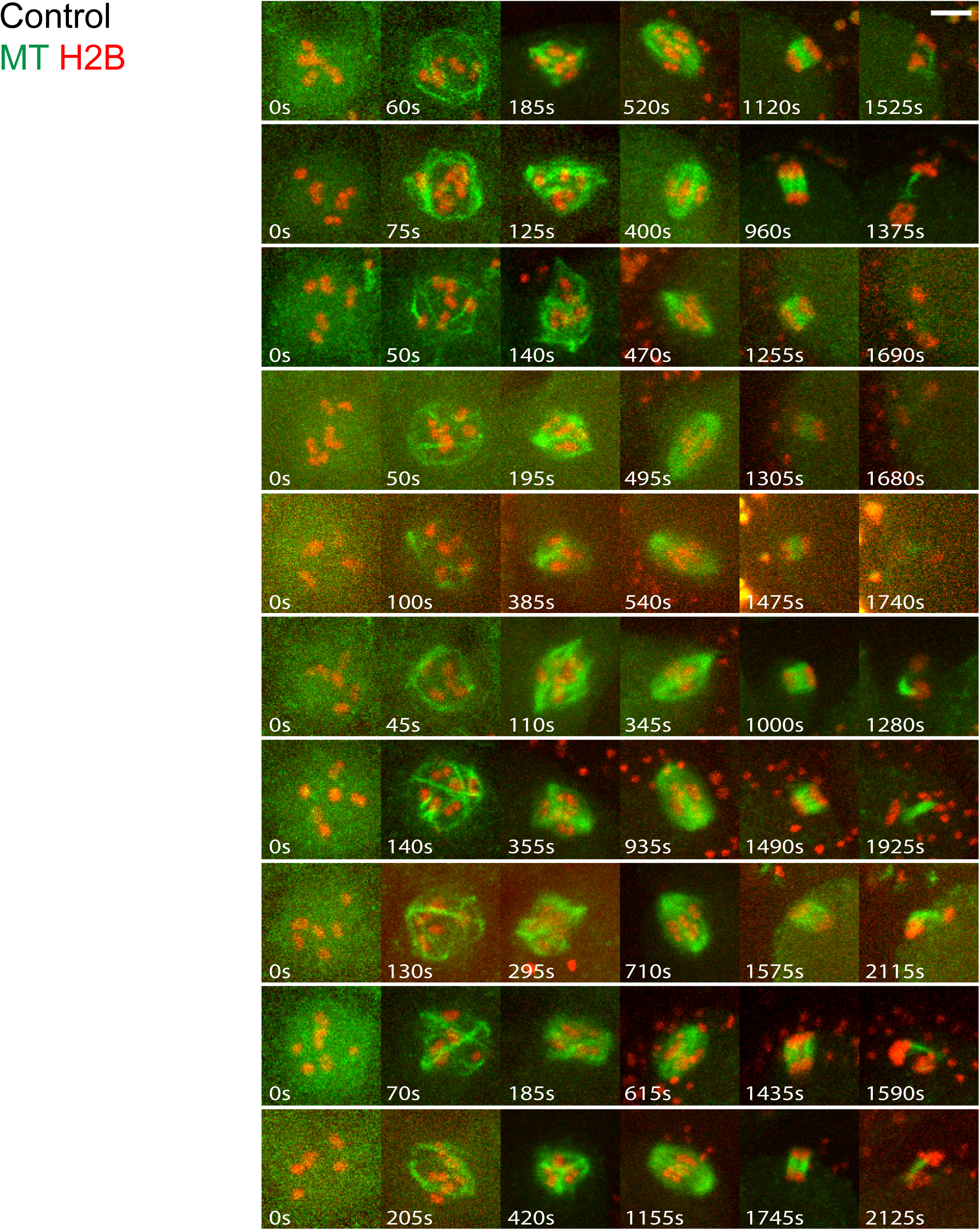
Time-lapse images during meiosis I for 10 live control oocytes expressing GFP::TBB-2 and mCherry::H2B.

**Supplemental Figure 2.**
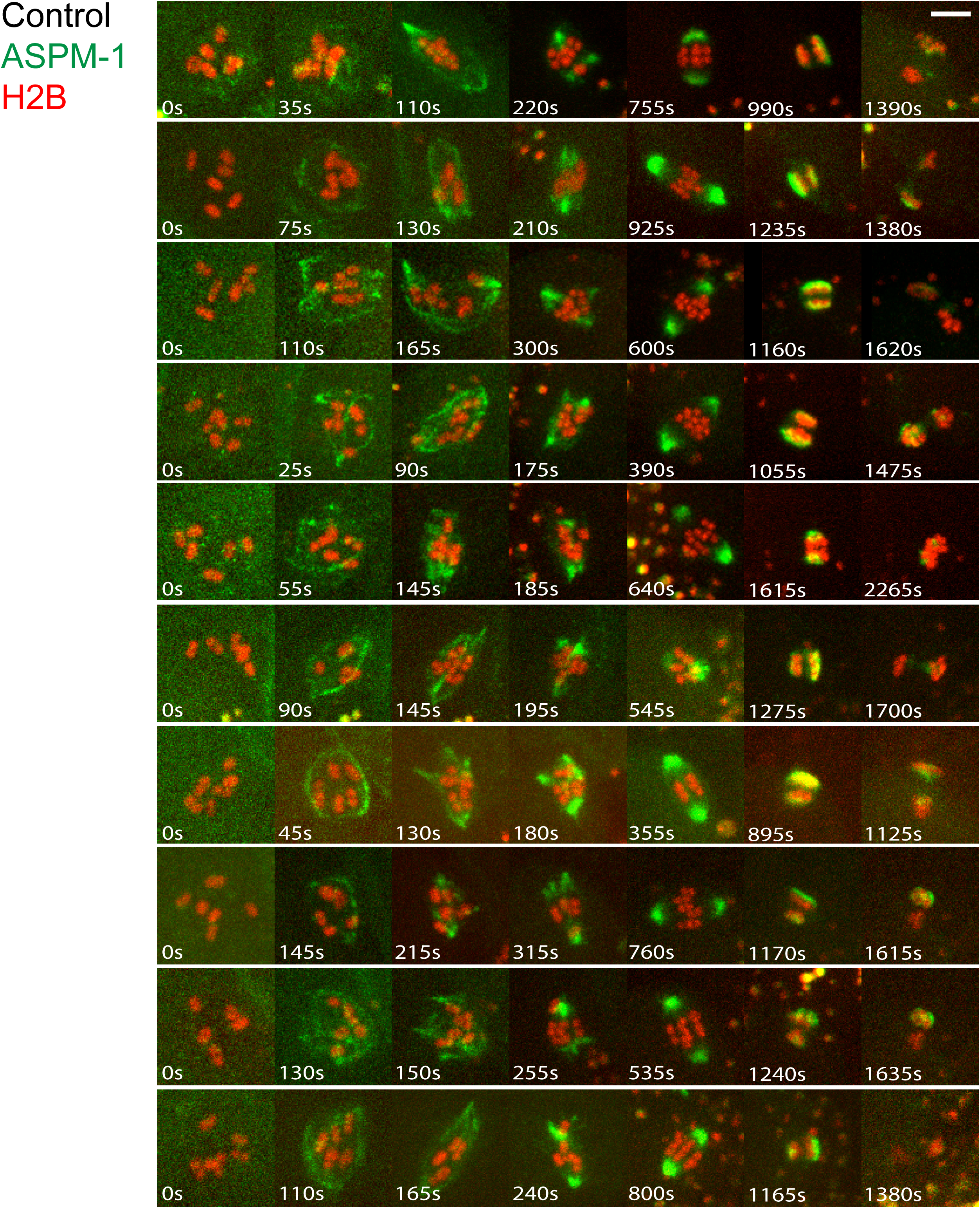
Time-lapse images during meiosis I for 10 live control oocytes expressing GFP::ASPM-1 and mCherry::H2B.

**Supplemental Figure 3.**
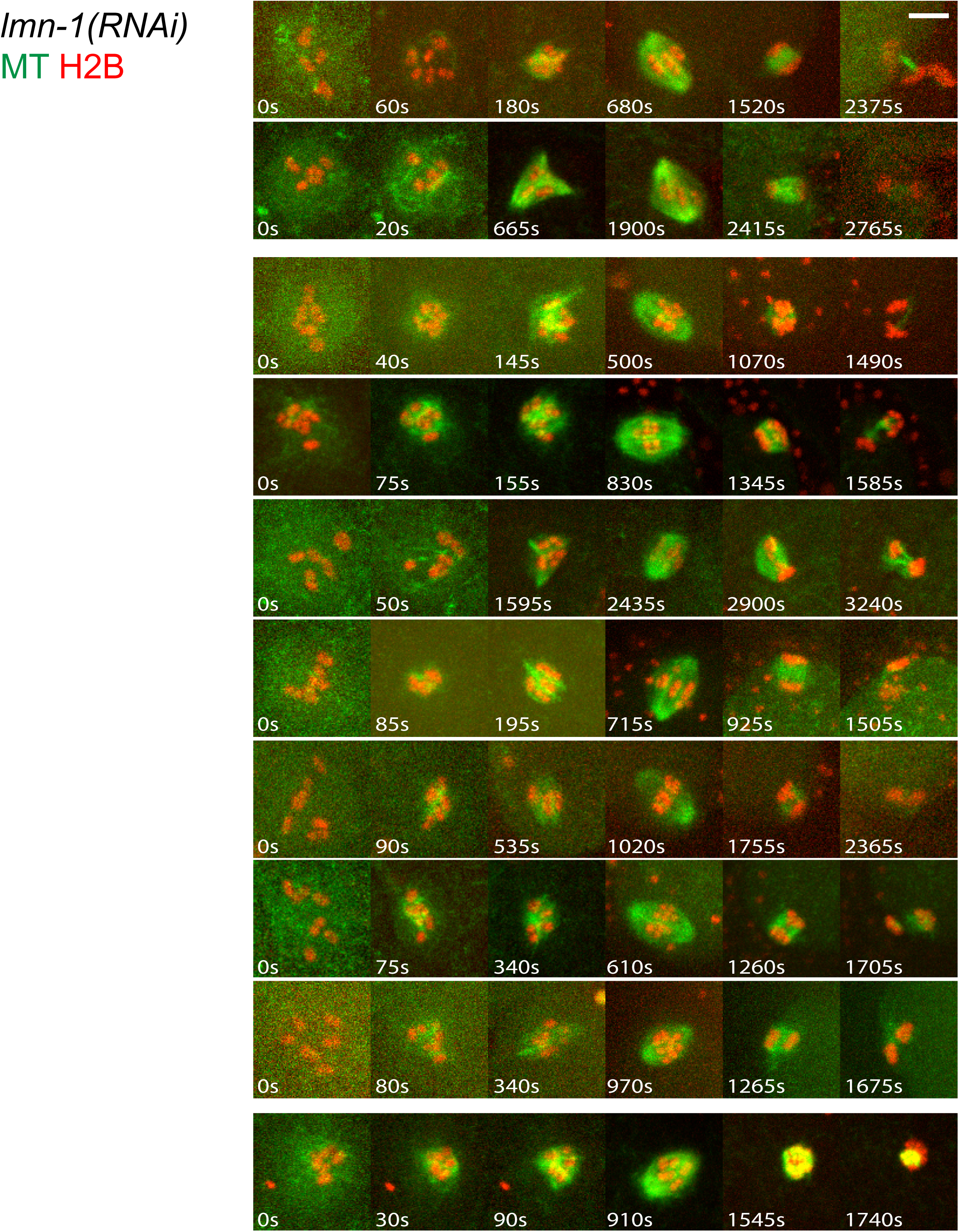
Time-lapse images during meiosis I for 10 live *lmn-1(RNAi)* oocytes expressing GFP::TBB-2 and mCherry::H2B. Rows 1 & 2: oocytes with cage structure; Rows 3-9 oocytes without cage structure; Row 10: oocyte without cage and no chromosome segregation. In all figures, cage structures and chromosome segregation we are assessed using Imaris software to rotate 3-D images.

**Supplemental Figure 4.**
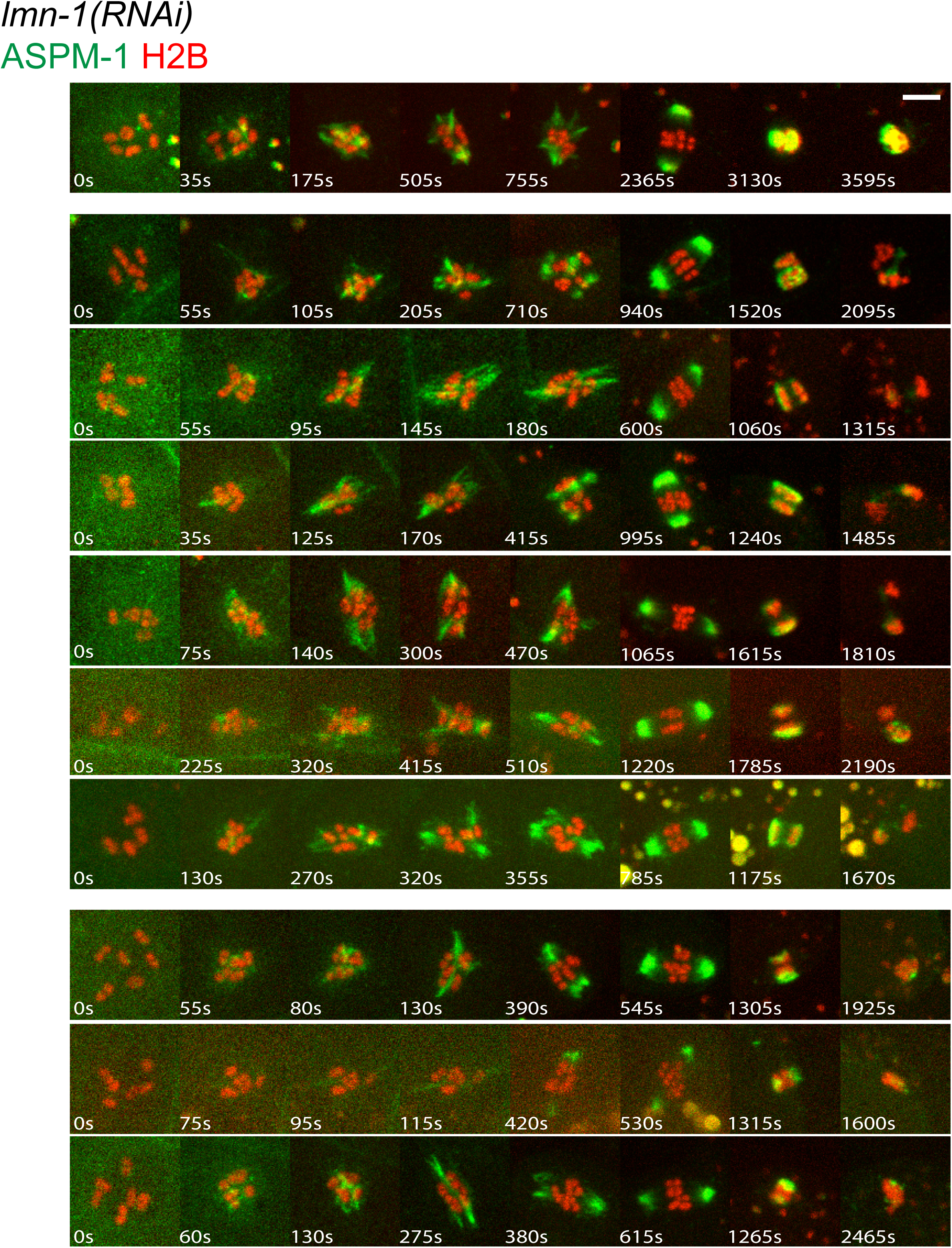
Time-lapse images during meiosis I for 10 live *lmn-1(RNAi)* oocytes expressing GFP::ASPM-1 and mCherry::H2B. Row 1: oocyte with cage structure and no chromosome segregation; Rows 2-7 oocytes without cage structure; Row 8-10: oocytes without cage and no chromosome segregation.

**Supplemental Figure 5.**
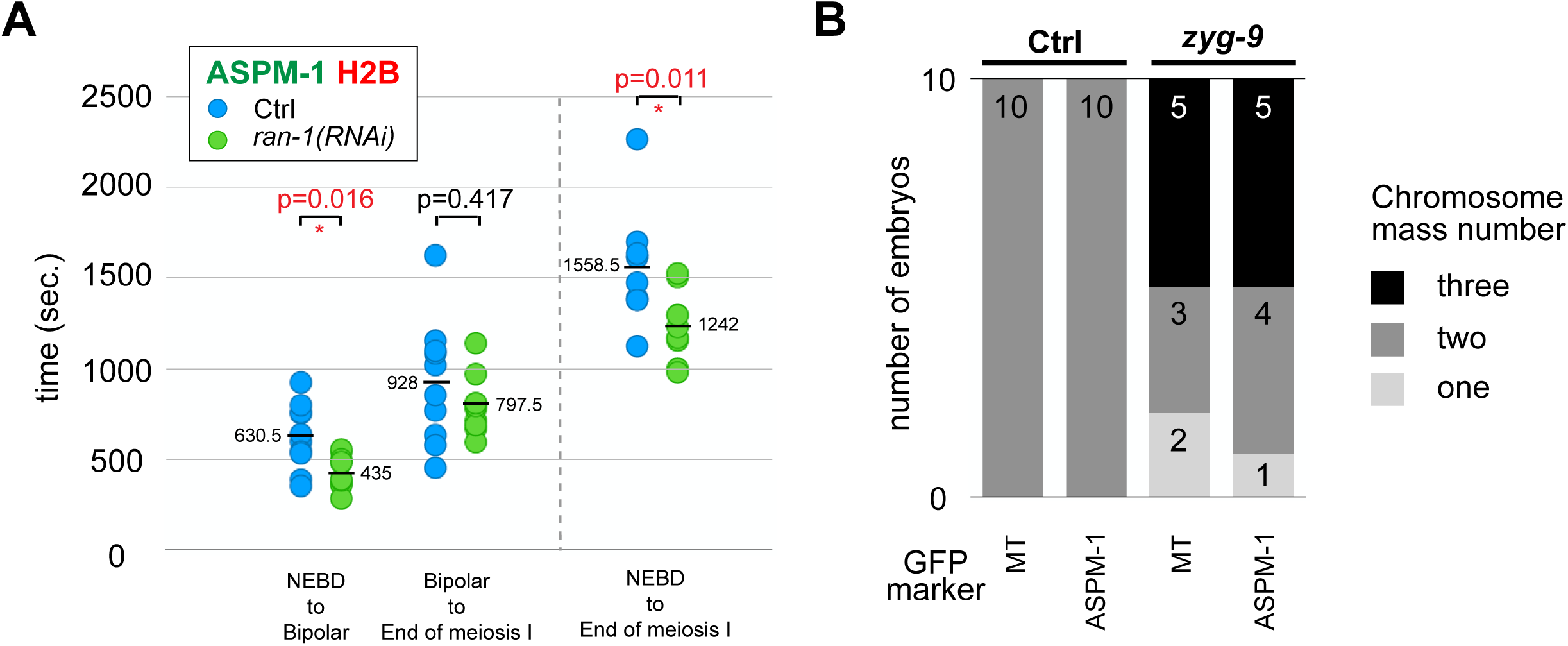
(A) Scatter plot showing comparisons of the time from NEBD to spindle bipolarity establishment (Bipolar), from spindle bipolarity establishment to the end of meiosis I, and entire time to progress through meiosis I between control and *ran-1(RNAi)* oocytes expressing GFP::ASPM-1 and mCherry::H2B. (B) The number of segregating chromosome masses detected in control and *zyg-9(RNAi)* oocytes at the end of meiosis I for strains expressing either GFP::TBB-2 and mCherry::H2B, or GFP::ASPM-1 and mCherry::H2B. Numbers within the bars indicate the number of embryos in each category.

**Supplemental Figure 6.**
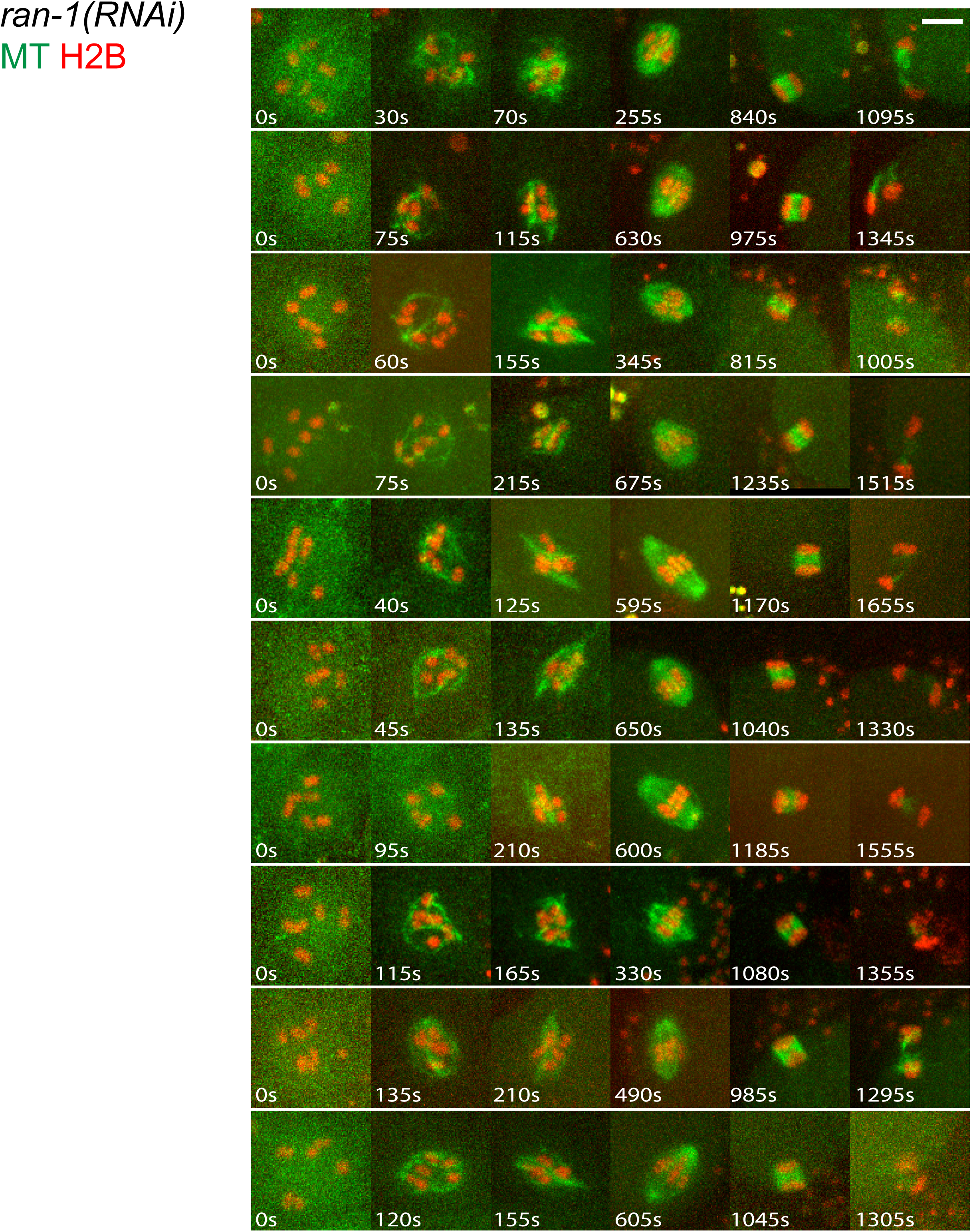
Time-lapse images during meiosis I for 10 live *ran-1(RNAi)* oocytes expressing GFP::TBB-2 and mCherry::H2B.

**Supplemental Figure 7.**
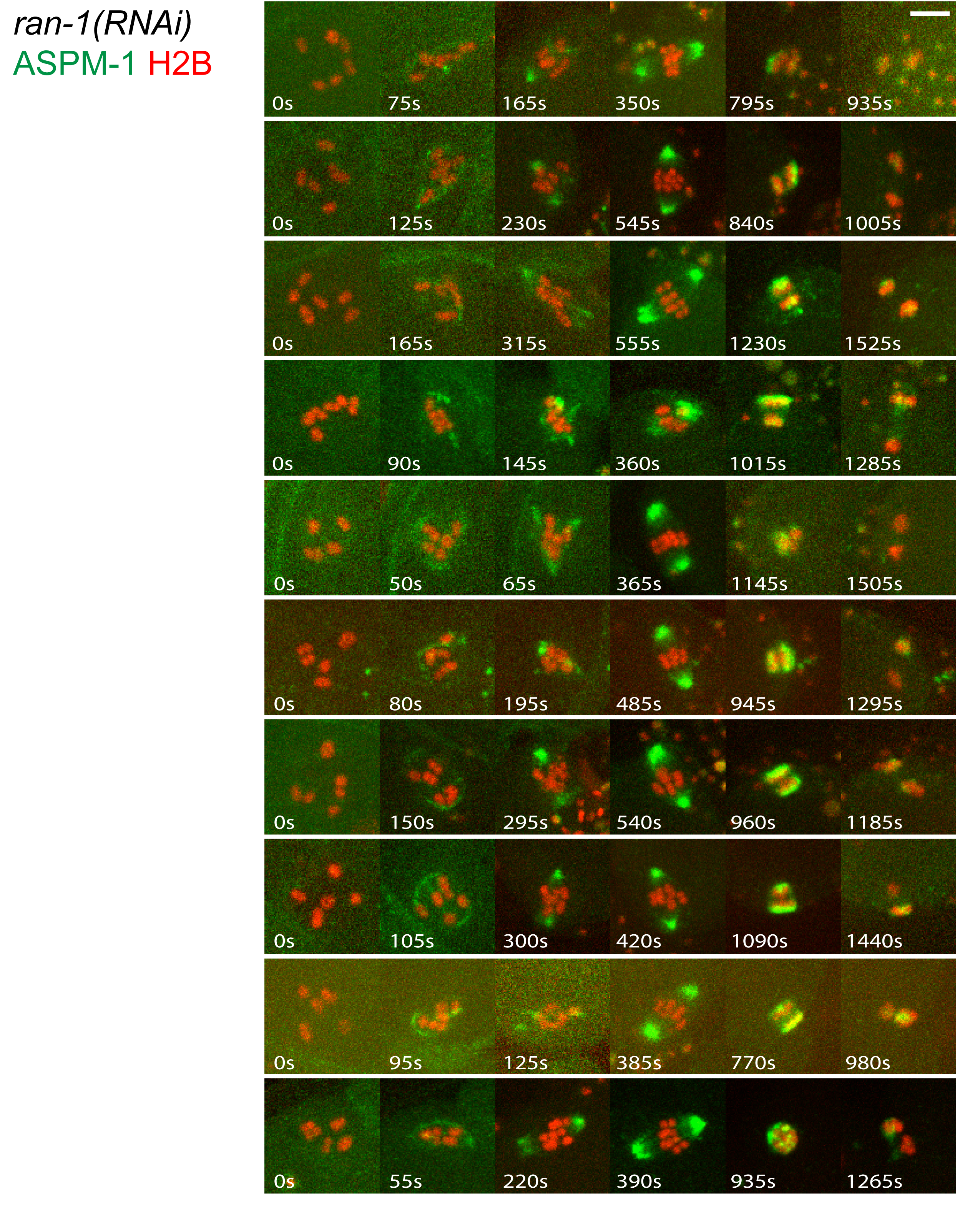
Time-lapse images during meiosis I for 10 live *ran-1(RNAi)* oocytes expressing GFP::ASPM-1 and mCherry::H2B.

**Supplemental Figure 8.**
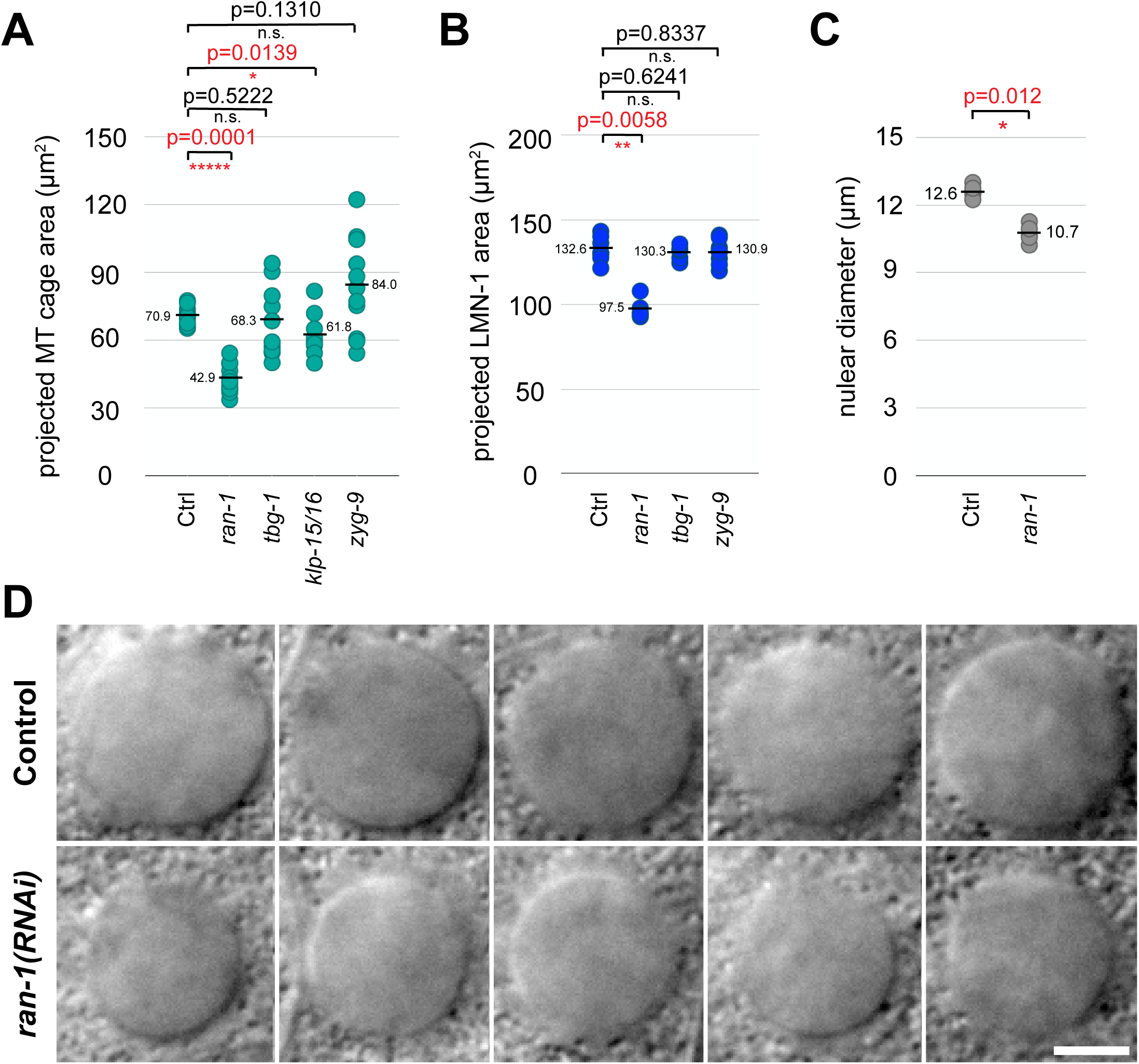
(A) Scatter plot showing projected areas of the microtubule cage-like structure meiosis I in control and mutant oocytes expressing GFP::TBB-2 and mCherry::H2B. (B) Scatter plot showing projected nuclear lamina area measured at the cage stage of meiosis I for control and mutant oocytes expressing GFP::LMN-1 and mCherry:: TBB-2. (C) Scatter plot showing nuclear diameter of control and *ran-1(RNAi)* oocytes measured using Nomarski images (8D). (D) Nomarski images of control and *ran-1(RNAi)* oocyte nuclei 2 seconds before nuclear envelope break down.

**Supplemental Figure 9.**
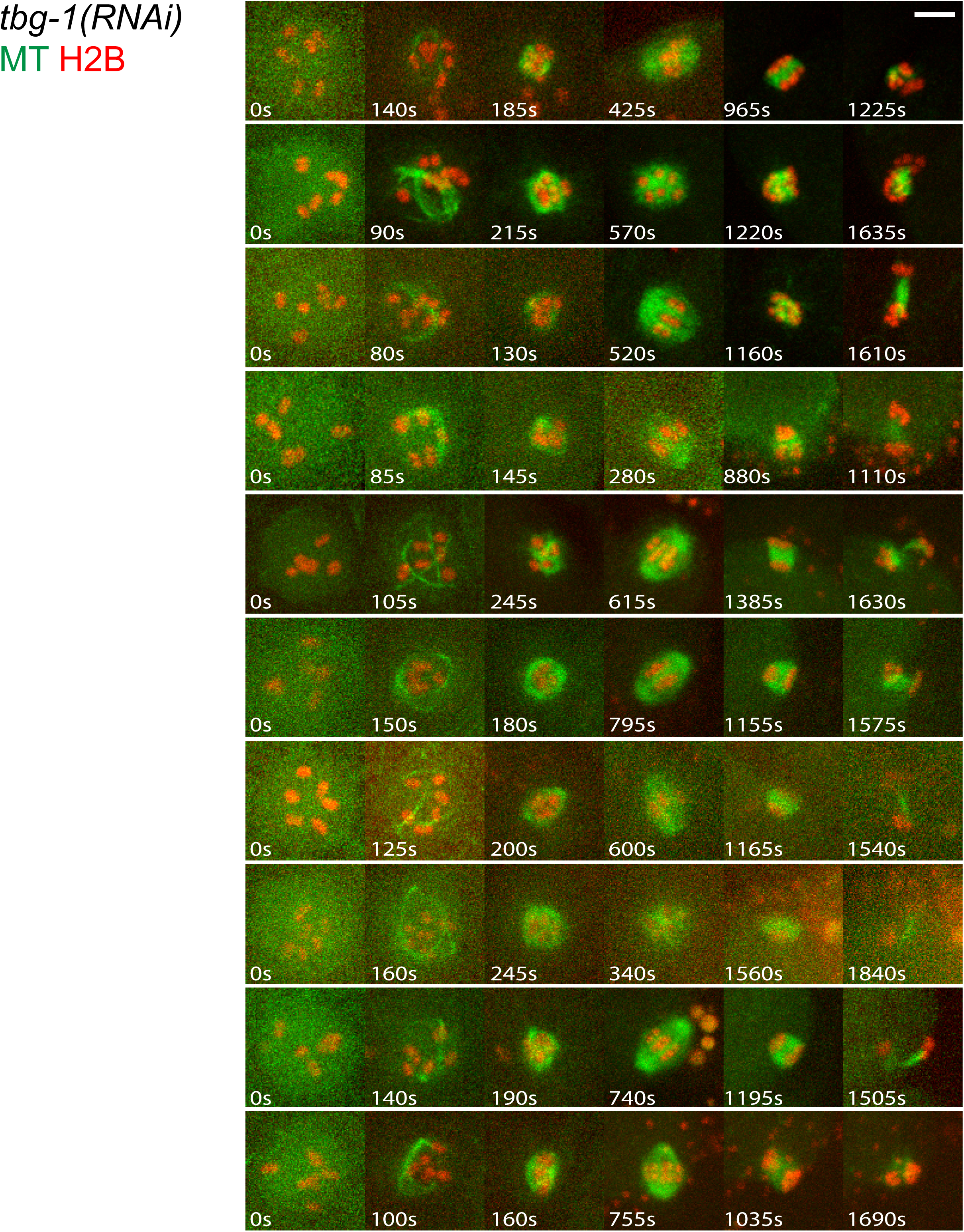
Time-lapse images during meiosis I for 10 live *tbg-1(RNAi)* oocytes expressing GFP::TBB-2 and mCherry::H2B.

**Supplemental Figure 10.**
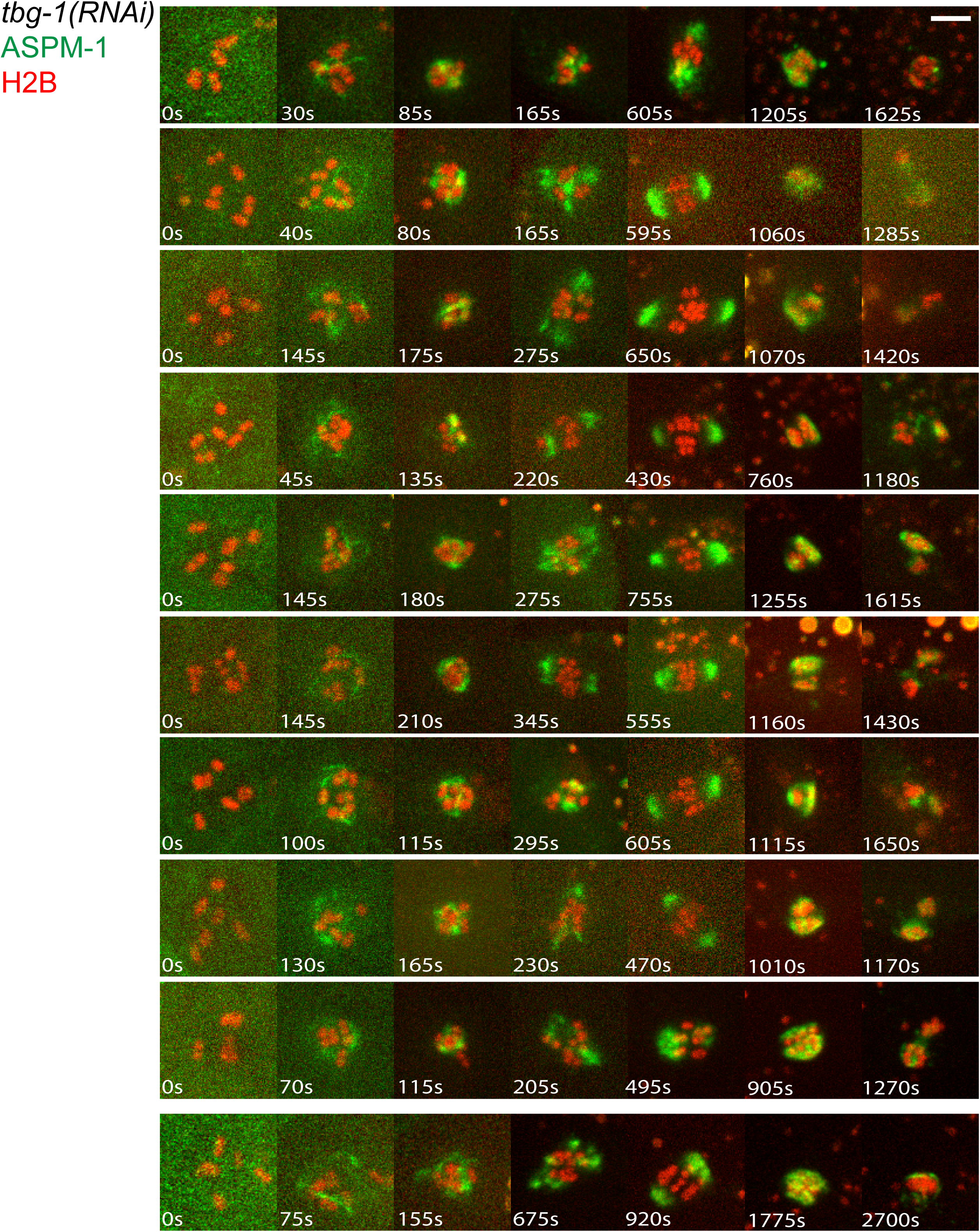
Time-lapse images during meiosis I for 10 live *tbg-1(RNAi)* oocytes expressing GFP::ASPM-1 and mCherry::H2B. Rows 1-9: oocytes with chromosome segregation; Row 10: oocyte without chromosome segregation.

**Supplemental Figure 11.**
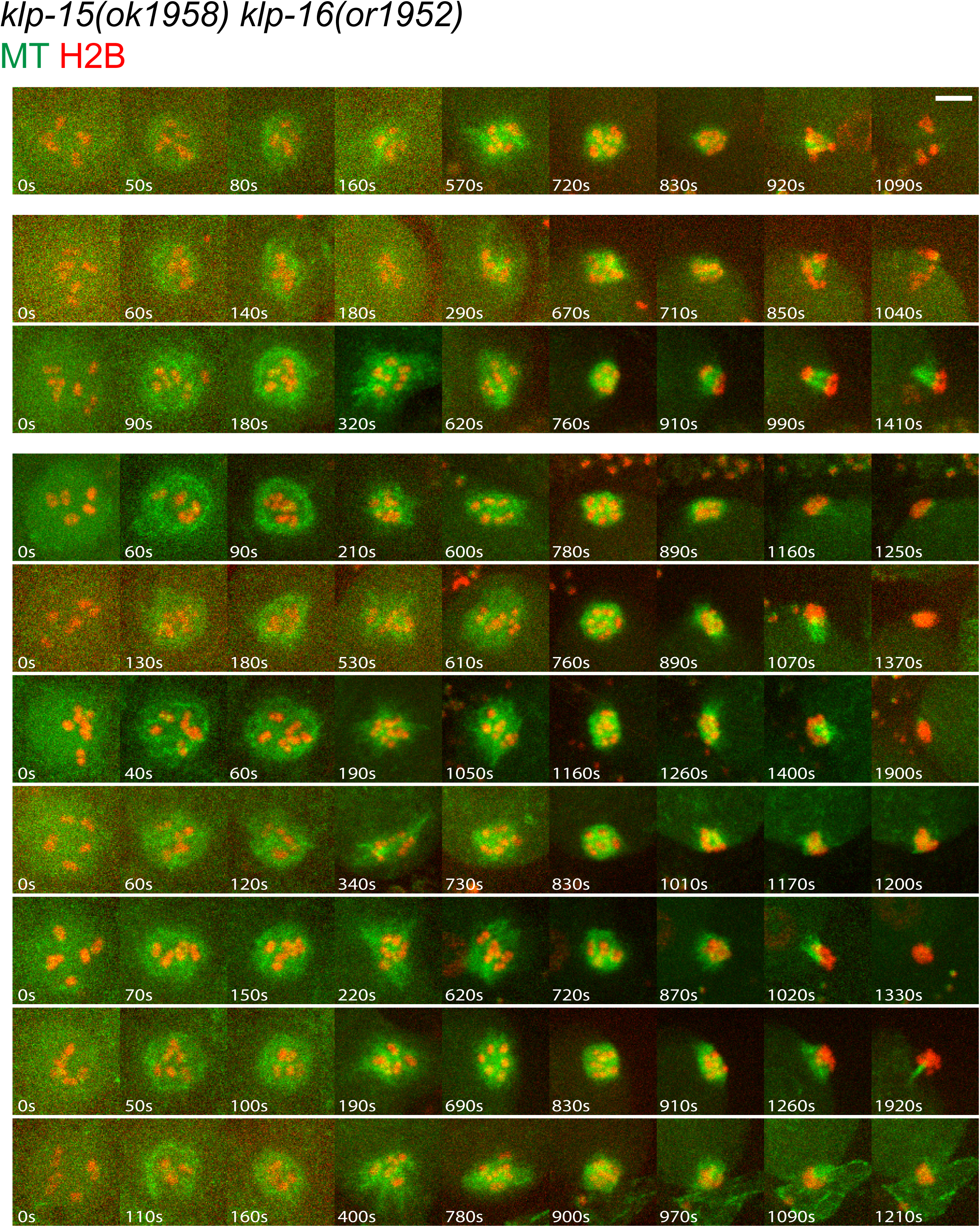
Time-lapse images during meiosis I for 10 live *klp-15/16(-/-)* double mutant oocytes expressing GFP::TBB-2 and mCherry::H2B. Row 1: oocyte with chromosomes segregated into three masses; Rows 2 & 3: oocytes with chromosomes segregated into two masses; Rows 4-10: oocytes with no chromosome segregation.

**Supplemental Figure 12.**
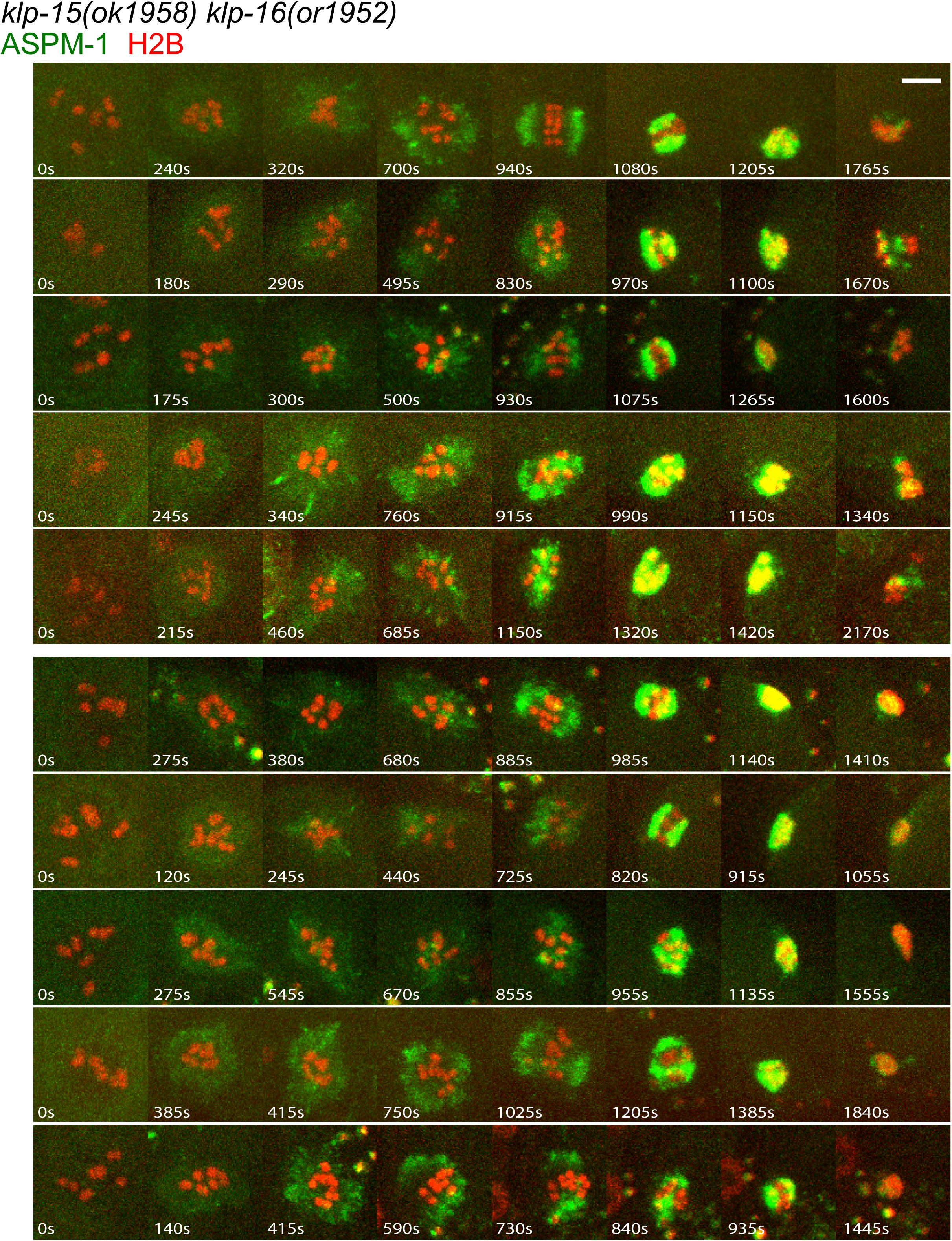
Time-lapse images during meiosis I for 10 live *klp-15/16(-/-)* double mutant oocytes expressing GFP::ASPM-1 and mCherry::H2B. Rows 1-5: oocytes with chromosomes segregated into two masses; Rows 6-10: oocytes with no chromosome segregation.

**Supplemental Figure 13.**
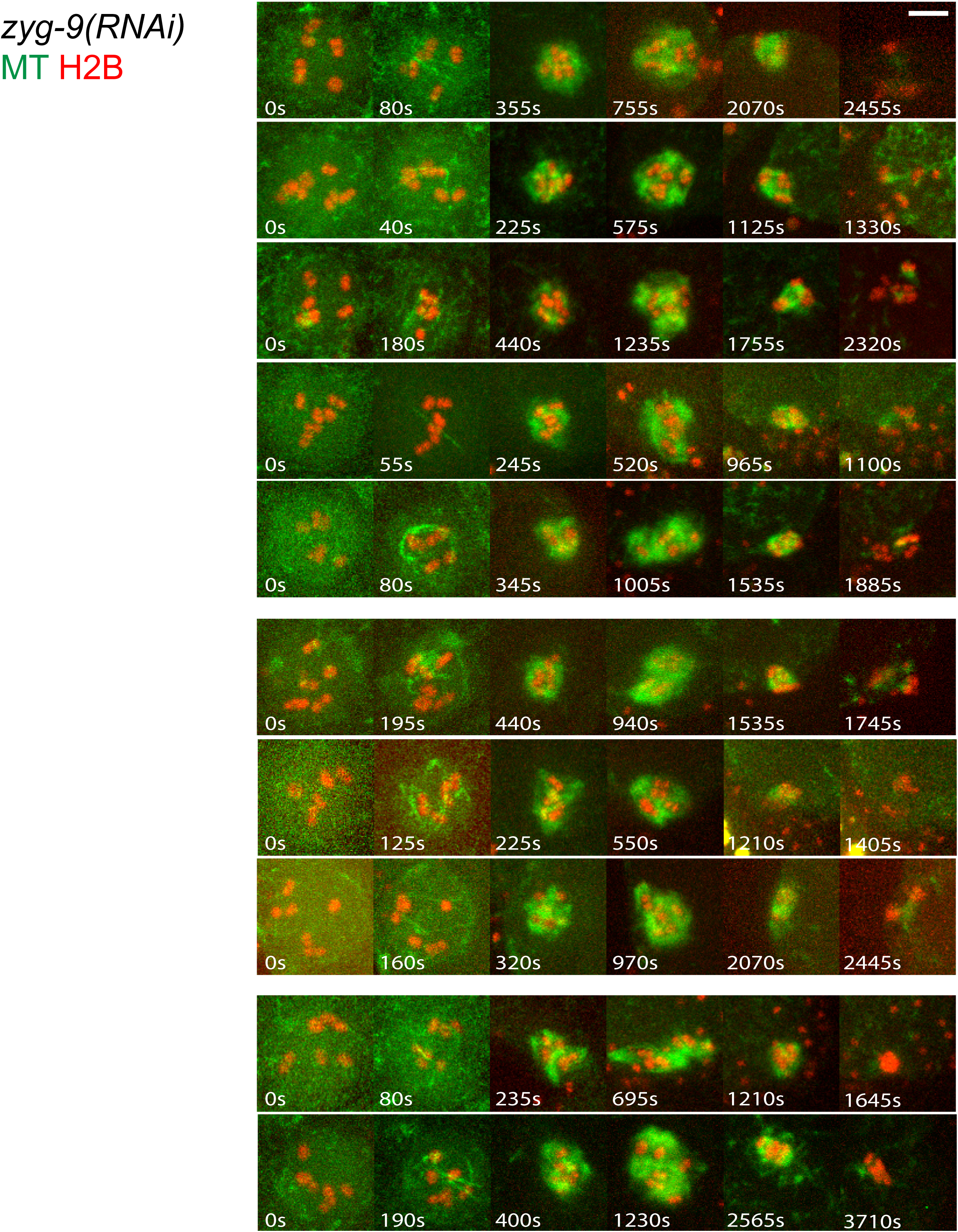
Time-lapse images during meiosis I for 10 live *zyg-9(RNAi)* oocytes expressing GFP::TBB-2 and mCherry::H2B. Rows 1-5: oocytes with chromosomes segregated into three masses; Rows 6-8: oocytes with chromosomes segregated into two masses; Rows 9 & 10: oocytes with no chromosome segregation.

**Supplemental Figure 14.**
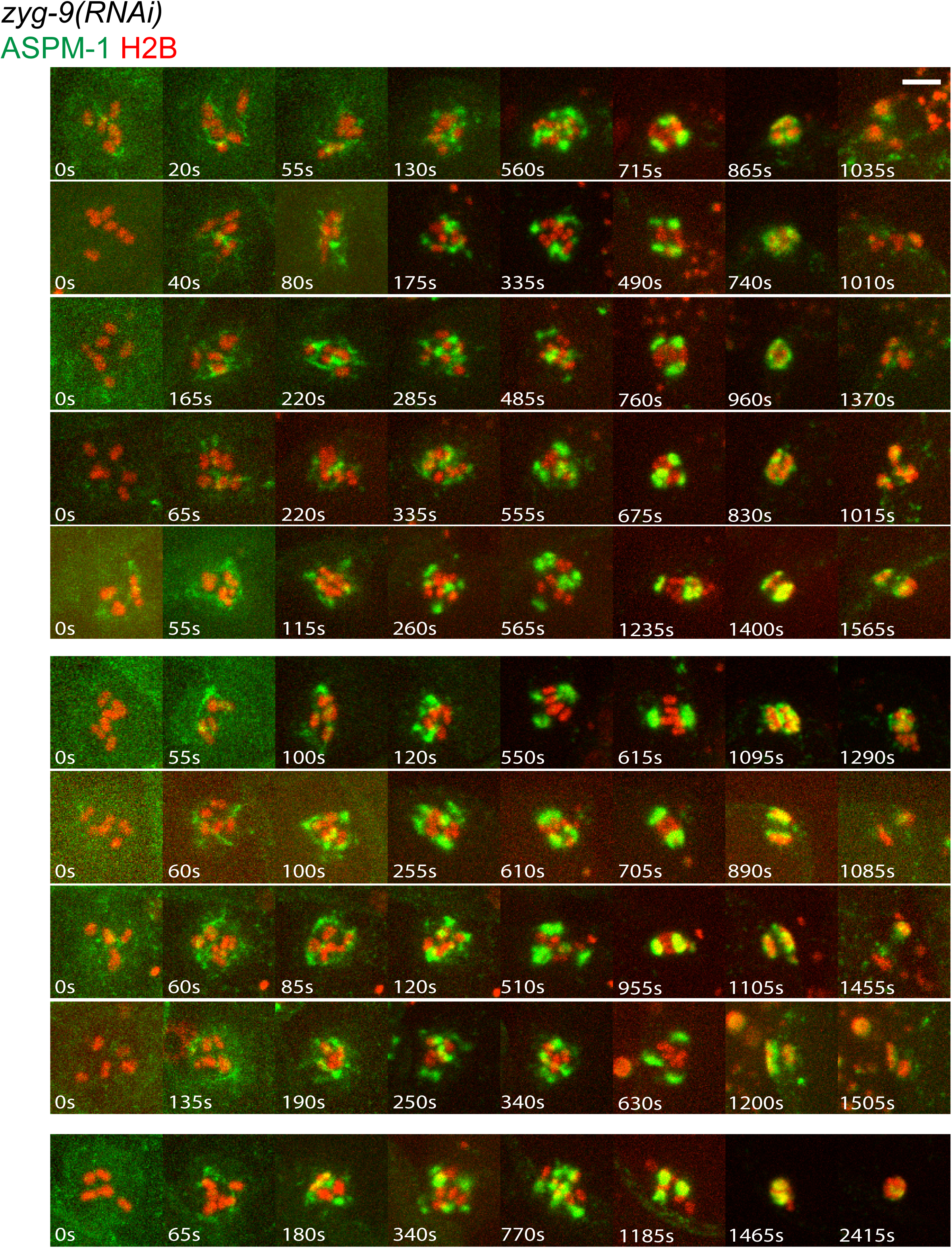
Time-lapse images during meiosis I for 10 live *zyg-9(RNAi)* oocytes expressing GFP::ASPM-1 and mCherry::H2B. Rows 1-5: oocytes with chromosomes segregated into 3 masses; Rows 6-9: oocytes with chromosomes segregated into two masses; Row 10: no chromosome segregation.

**Supplemental Figure 15.**
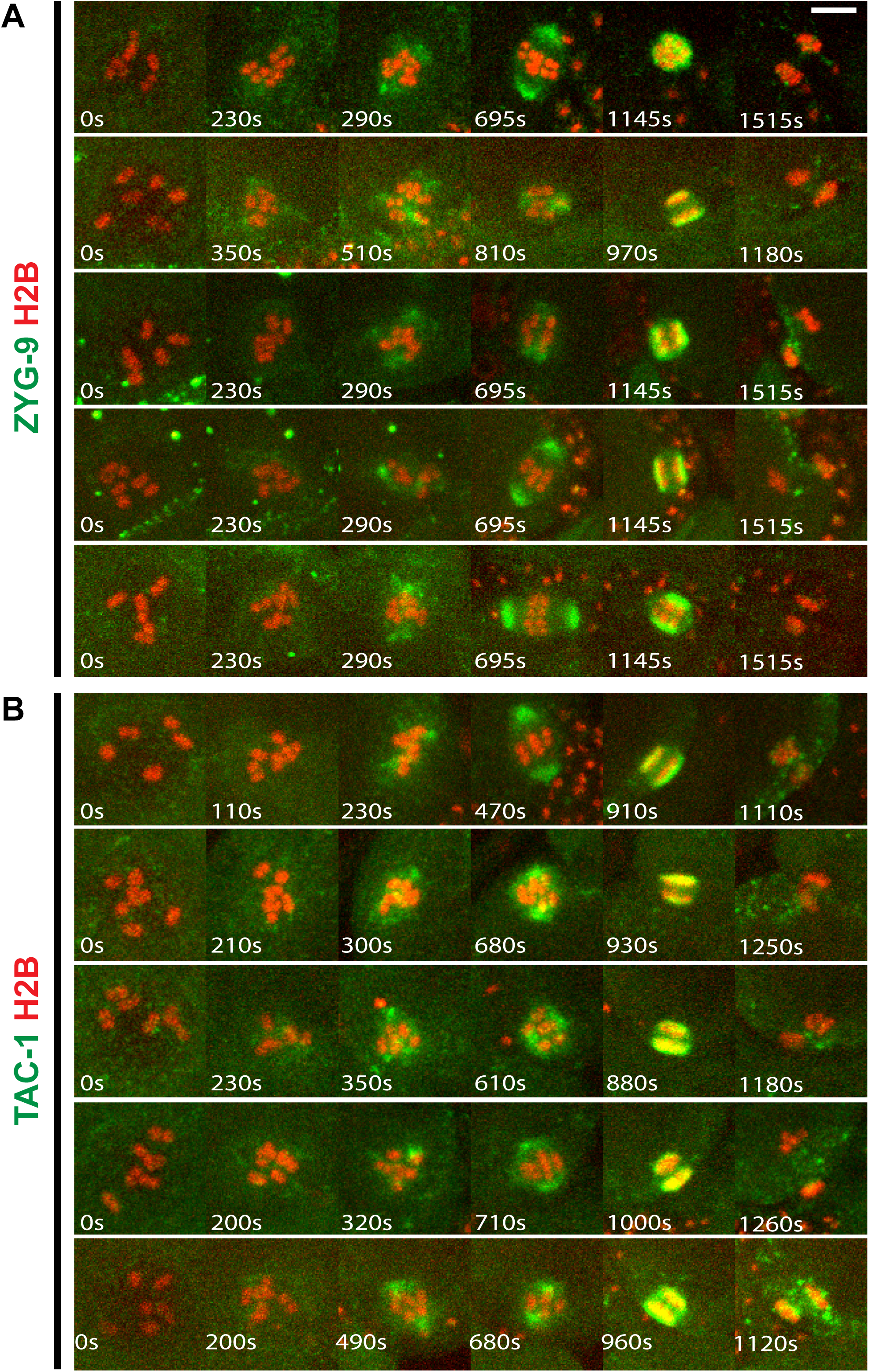
Time-lapse images during meiosis I for live control oocytes expressing either GFP::ZYG-9 and mCherry::H2B (A), or GFP::TAC-1 and mCherry::H2B (B).

**Supplemental Figure 16.**
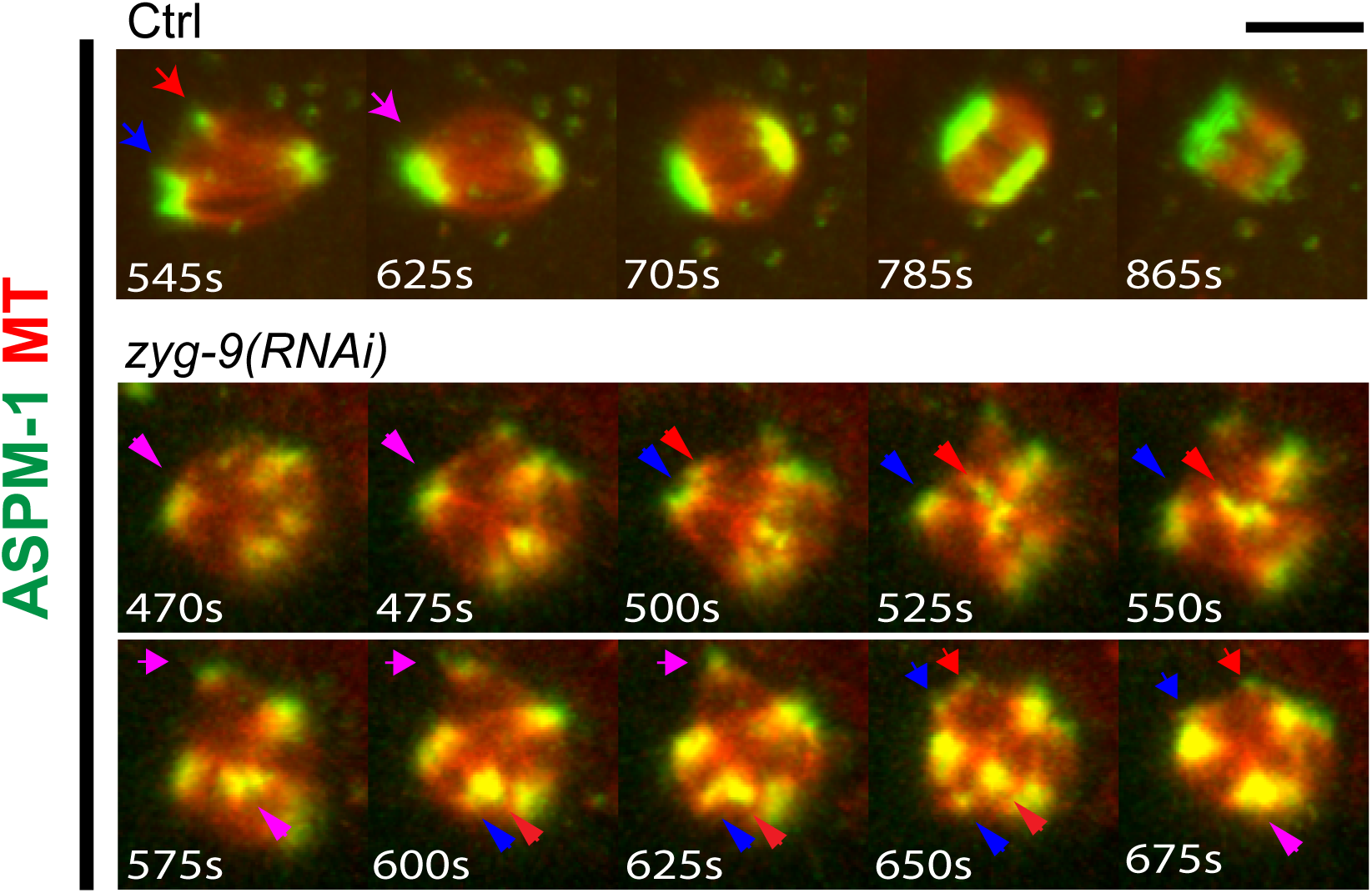
Time-lapse images during meiosis I for live control and *zyg-9(RNAi)* oocytes expressing GFP::ASPM-1 and mCherry:: TBB-2. Times after NEBD are indicated. Pink arrowheads indicate spindle poles that split into two foci that are then marked by red and blue arrowheads.

**Supplemental Table 1.**

| Strain name | Genotype |
| --- | --- |
| N2 | (ancestral) |
| EU2876 | *aspm-1(or1935[gfp::aspm-1]) I; itIs37[pie-1p::mCherry::H2B::pie-1 3'UTR + unc-119(+)] IV* |
| EU2942 | *ruls57[pie-1p::gfp::tubulin + unc-119(+)]; itIs37[pie-1p::mCherry::H2B::pie-1 3'UTR + unc-119(+)] IV* |
| EU3006 | *aspm-1(or1935[gfp::aspm-1]) I; tbb-2(tj29[mCherry::tbb-2]) III* |
| EU3091 | *ijmSi49[pJD479; Pmex-5::gfp::lmn-1::3'tbb-2] I; tbb-2(tj29[mCherry::tbb-2]) III* |
| EU3115 | *klp-15(ok1958) klp-16(or1952)/ tmC18[dpy-5(tmIs1236)] I; itIs37[pie-1p::mCherry::H2B::pie-1 3'UTR + unc-119(+)] IV; ruls57[pie-1p::gfp::tubulin + unc-119(+)]* |
| EU3121 | *tac-1(or1955[gfp::tac-1]) II; itIs37[pie-1p::mCherry::H2B::pie-1 3'UTR + unc-119(+)] IV (may contain unc-119(ed3) III)* |
| EU3169 | *zyg-9(or1956[gfp::zyg-9]) II; itIs37[pie-1p::mCherry::H2B, unc-119(+)] IV (may contain unc-119(ed3) III)* |
| RB1593 | *klp-15(ok1958)* |
| EU3201 | *klp-15(ok1958) aspm-1(syb1260[gfp::aspm-1]) klp-16(or1952) /tmC18[dpy-5(tmIs1236)] I; itIs37[pie-1p::mCherry::H2B::pie-1 3'UTR + unc-119(+)] IV* |

**Supplemental Movie 1.** Time-lapse movie during meiosis I of a live control oocyte expressing GFP::ASPM-1 and mCherry::H2B.

**Supplemental Movies 2 and 3.** Time-lapse movies during meiosis I for live *klp-15/16(-/-)* double mutant oocytes expressing GFP::ASPM-1 and mCherry::H2B.

**Supplemental Movies 4, 5 and 6.** Three-dimensional rotation movies during meiosis I for live control oocytes expressing GFP::TBB-2 and mCherry::H2B.

**Supplemental Movies 7, 8 and 9.** Three-dimensional rotation movies during meiosis I for live *zyg-9(RNAi)* oocytes expressing GFP::TBB-2 and mCherry::H2B.

**Supplemental Movie 10.** *Ex utero* time-lapse spinning disk confocal maximum intensity z projection of 15 planes with 1 μm z-spacing movies during meiosis I for live control oocyte expressing GFP::TBB-2 and mCherry::H2B.

**Supplemental Movie 11.** *Ex utero*, time-lapse spinning disk confocal maximum intensity z-projection for 15 planes with 1 μm z-spacing movies during meiosis I of live *zyg-9(RNAi)* oocyte expressing GFP::TBB-2 and mCherry::H2B.

**Supplemental Movie 12** Time-lapse movie during meiosis I for live control oocyte expressing GFP::ASPM-1 and mCherry:: TBB-2.

**Supplemental Movies 13 and 14.** Time-lapse movies during meiosis I for live *zyg-9(RNAi)* oocyte expressing GFP::ASPM-1 and mCherry::H2B.

**Supplemental Movie 15** Time-lapse movies during meiosis I for live *zyg-9(RNAi)* oocyte expressing GFP::ASPM-1 and mCherry::TBB-2.

**Supplemental Movie 16** Three-dimensional rotation movie for live *tac-1(RNAi)* oocyte expressing GFP::TBB-2 and mCherry::H2B.

**Supplemental Movie 17** Time-lapse movies during meiosis I for live *tac-1(RNAi)* oocyte expressing GFP::ASPM-1 and mCherry::H2B.

## REFERENCES

Akhmanova, A. and Steinmetz, M.O., (2015). Control of microtubule organization and dynamics: two ends in the limelight. Nat Rev Mol Cell Biol 16, 711–726.

Askjaer, P., Galy, V., Hannak, E. and Mattaj, I.W., (2002). Ran GTPase cycle and importins alpha and beta are essential for spindle formation and nuclear envelope assembly in living Caenorhabditis elegans embryos. Mol Biol Cell 13, 4355–4370.

Bellanger, J.M., Carter, J.C., Phillips, J.B., Canard, C., Bowerman, B. and Gonczy, P., (2007). ZYG-9, TAC-1 and ZYG-8 together ensure correct microtubule function throughout the cell cycle of C. elegans embryos. J Cell Sci 120, 2963–2973.

Bellanger, J.M. and Gonczy, P., (2003). TAC-1 and ZYG-9 form a complex that promotes microtubule assembly in C. elegans embryos. Curr Biol 13, 1488–1498.

Brittle, A.L. and Ohkura, H., (2005). Mini spindles, the XMAP215 homologue, suppresses pausing of interphase microtubules in Drosophila. EMBO J 24, 1387–1396.

Clarke, P.R. and Zhang, C., (2008). Spatial and temporal coordination of mitosis by Ran GTPase. Nat Rev Mol Cell Biol 9, 464–477.

Connolly, A.A., Sugioka, K., Chuang, C.-H., Lowry, J.B. and Bowerman, B., (2015). KLP-7 acts through the Ndc80 complex to limit pole number in C. elegans oocyte meiotic spindle assembly. J Cell Biol 210, 917–932.

Cullen, C.F. and Ohkura, H., (2001). Msps protein is localized to acentrosomal poles to ensure bipolarity of Drosophila meiotic spindles. Nat Cell Biol 3, 637–642.

Dejima, K., Hori, S., Iwata, S., Suehiro, Y., Yoshina, S., Motohashi, T. and Mitani, S., (2018). An Aneuploidy-Free and Structurally Defined Balancer Chromosome Toolkit for Caenorhabditis elegans. Cell Rep 22, 232–241.

Dumont, J. and Desai, A., (2012). Acentrosomal spindle assembly and chromosome segregation during oocyte meiosis. Trends Cell Biol 22, 241–249.

Edzuka, T., Yamada, L., Kanamaru, K., Sawada, H. and Goshima, G., (2014). Identification of the augmin complex in the filamentous fungus Aspergillus nidulans. PLoS One 9, e101471.

Fink, G., Hajdo, L., Skowronek, K.J., Reuther, C., Kasprzak, A.A. and Diez, S., (2009). The mitotic kinesin-14 Ncd drives directional microtubule-microtubule sliding. Nat Cell Biol 11, 717–723.

Gigant, E., Stefanutti, M., Laband, K., Gluszek-Kustusz, A., Edwards, F., Lacroix, B., Maton, G., Canman, J.C., Welburn, J.P.I. and Dumont, J., (2017). Inhibition of ectopic microtubule assembly by the kinesin-13 KLP-7 prevents chromosome segregation and cytokinesis defects in oocytes. Development 144, 1674–1686.

Goodman, B., Channels, W., Qiu, M., Iglesias, P., Yang, G. and Zheng, Y., (2010). Lamin B counteracts the kinesin Eg5 to restrain spindle pole separation during spindle assembly. J Biol Chem 285, 35238–35244.

Goshima, G., Nedelec, F. and Vale, R.D., (2005). Mechanisms for focusing mitotic spindle poles by minus end-directed motor proteins. J Cell Biol 171, 229–240.

Gunzelmann, J., Ruthnick, D., Lin, T.C., Zhang, W., Neuner, A., Jakle, U. and Schiebel, E., (2018). The microtubule polymerase Stu2 promotes oligomerization of the gamma-TuSC for cytoplasmic microtubule nucleation. Elife 7.

Hatsumi, M. and Endow, S.A., (1992). The Drosophila ncd microtubule motor protein is spindle-associated in meiotic and mitotic cells. J Cell Sci 103 (Pt 4), 1013–1020.

Hayashi, D., Tanabe, K., Katsube, H. and Inoue, Y.H., (2016). B-type nuclear lamin and the nuclear pore complex Nup107-160 influences maintenance of the spindle envelope required for cytokinesis in Drosophila male meiosis. Biol Open 5, 1011–1021.

Ito, A. and Goshima, G., (2015). Microcephaly protein Asp focuses the minus ends of spindle microtubules at the pole and within the spindle. J Cell Biol 211, 999–1009.

Kamath, R.S., Fraser, A.G., Dong, Y., Poulin, G., Durbin, R., Gotta, M., Kanapin, A., Le Bot, N., Moreno, S., Sohrmann, M., Welchman, D.P., Zipperlen, P. and Ahringer, J., (2003). Systematic functional analysis of the Caenorhabditis elegans genome using RNAi. Nature 421, 231–237.

Kamath, R.S., Martinez-Campos, M., Zipperlen, P., Fraser, A.G. and Ahringer, J., (2001). Effectiveness of specific RNA-mediated interference through ingested double-stranded RNA in Caenorhabditis elegans. Genome Biol 2, RESEARCH0002.

Kosco, K.A., Pearson, C.G., Maddox, P.S., Wang, P.J., Adams, I.R., Salmon, E.D., Bloom, K. and Huffaker, T.C., (2001). Control of microtubule dynamics by Stu2p is essential for spindle orientation and metaphase chromosome alignment in yeast. Mol Biol Cell 12, 2870–2880.

Liu, J., Rolef Ben-Shahar, T., Riemer, D., Treinin, M., Spann, P., Weber, K., Fire, A. and Gruenbaum, Y., (2000). Essential roles for Caenorhabditis elegans lamin gene in nuclear organization, cell cycle progression, and spatial organization of nuclear pore complexes. Mol Biol Cell 11, 3937–3947.

Matthies, H.J., McDonald, H.B., Goldstein, L.S. and Theurkauf, W.E., (1996). Anastral meiotic spindle morphogenesis: role of the non-claret disjunctional kinesin-like protein. The Journal of Cell Biology 134, 455–464.

McNally, K., Audhya, A., Oegema, K. and McNally, F.J., (2006). Katanin controls mitotic and meiotic spindle length. J Cell Biol 175, 881–891.

Mullen, T.J., Davis-Roca, A.C. and Wignall, S.M., (2019). Spindle assembly and chromosome dynamics during oocyte meiosis. Curr Opin Cell Biol 60, 53–59.

Mullen, T.J. and Wignall, S.M., (2017). Interplay between microtubule bundling and sorting factors ensures acentriolar spindle stability during C. elegans oocyte meiosis. PLoS Genet 13, e1006986.

Ohkura, H., (2015). Meiosis: an overview of key differences from mitosis. Cold Spring Harb Perspect Biol 7.

Schuh, M. and Ellenberg, J., (2007). Self-organization of MTOCs replaces centrosome function during acentrosomal spindle assembly in live mouse oocytes. Cell 130, 484–498.

Severson, A.F., von Dassow, G. and Bowerman, B., (2016). Oocyte Meiotic Spindle Assembly and Function. Curr Top Dev Biol 116, 65–98.

Shirasu-Hiza, M., Coughlin, P. and Mitchison, T., (2003). Identification of XMAP215 as a microtubule-destabilizing factor in Xenopus egg extract by biochemical purification. J Cell Biol 161, 349–358.

Skold, H.N., Komma, D.J. and Endow, S.A., (2005). Assembly pathway of the anastral Drosophila oocyte meiosis I spindle. J Cell Sci 118, 1745–1755.

So, C., Seres, K.B., Steyer, A.M., Monnich, E., Clift, D., Pejkovska, A., Mobius, W. and Schuh, M., (2019). A liquid-like spindle domain promotes acentrosomal spindle assembly in mammalian oocytes. Science 364.

Srayko, M., O’Toole E, T., Hyman, A.A. and Müller-Reichert, T., (2006). Katanin disrupts the microtubule lattice and increases polymer number in C. elegans meiosis. Curr Biol 16, 1944–1949.

Thawani, A., Kadzik, R.S. and Petry, S., (2018). XMAP215 is a microtubule nucleation factor that functions synergistically with the gamma-tubulin ring complex. Nat Cell Biol 20, 575–585.

Timmons, L. and Fire, A., (1998). Specific interference by ingested dsRNA. Nature 395, 854.

Tsai, M.Y., Wang, S., Heidinger, J.M., Shumaker, D.K., Adam, S.A., Goldman, R.D. and Zheng, Y., (2006). A mitotic lamin B matrix induced by RanGTP required for spindle assembly. Science 311, 1887–1893.

Woodruff, J.B., Ferreira Gomes, B., Widlund, P.O., Mahamid, J., Honigmann, A. and Hyman, A.A., (2017). The Centrosome Is a Selective Condensate that Nucleates Microtubules by Concentrating Tubulin. Cell 169, 1066–1077 e1010.

Yang, H.Y., McNally, K. and McNally, F.J., (2003). MEI-1/katanin is required for translocation of the meiosis I spindle to the oocyte cortex in C elegans. Dev Biol 260, 245–259.

Zhang, D., Tu, S., Stubna, M., Wu, W.S., Huang, W.C., Weng, Z. and Lee, H.C., (2018). The piRNA targeting rules and the resistance to piRNA silencing in endogenous genes. Science 359, 587–592.

Zhang, L., Ward, J.D., Cheng, Z. and Dernburg, A.F., (2015). The auxin-inducible degradation (AID) system enables versatile conditional protein depletion in C. elegans. Development 142, 4374–4384.

